# Depletion of microenvironmental syndecan-2 impairs hematopoietic stem cell self-renewal and cytokine responses

**DOI:** 10.1101/2025.11.07.687077

**Authors:** Matthew W. Hagen, Nicollette J. Setiawan, Rachel Wellington, Kelsey A. Woodruff, Carli Newman, Taylor M. Billings, Mary N. Nazzaro, Brandon Hadland, Christina M. Termini

## Abstract

Syndecan-2 is a heparan sulfate proteoglycan highly enriched on murine bone marrow hematopoietic stem cells (HSCs) compared to terminally differentiated hematopoietic cells. Syndecan-2 binds growth factors via its heparan sulfate glycosaminoglycan chains to coordinate cell signaling. Knockdown of syndecan-2 reduces HSC self-renewal ability and promotes cell cycling via *Cdkn1c*. In this study, we analyzed the function of syndecan-2 expressed by bone marrow niche cells in hematopoiesis and HSC self-renewal. We determined that syndecan-2 is highly expressed by bone marrow mesenchymal stromal cells and moderately expressed by endothelial cells. To test the function of niche-expressed syndecan-2 in hematopoiesis, we generated transgenic mice depleted of *Sdc2* in *Lepr-*targeted mesenchymal stromal cells (*Sdc2*^ΔMSC^ mice) or *Cdh5*-targeted endothelial cells (*Sdc2*^ΔEC^ mice). Loss of syndecan-2 from endothelial or mesenchymal stromal cells did not change bone marrow HSC frequencies or numbers. However, depletion of syndecan-2 from *Lepr*-targeted mesenchymal stromal cells, but not *Cdh5*-targeted endothelial cells, diminishes HSC self-renewal ability analyzed by competitive transplants into lethally irradiated mice. *Ex vivo* studies further show that HSCs co-cultured with HS-5 stromal cells depleted of *SDC2* exhaust more rapidly than HSCs cultured with control HS-5 cells. Single-cell RNA sequencing analyses reveal that the depletion of *Sdc2* from mesenchymal stromal cells significantly remodels the HSC transcriptome by enriching for pathways associated with excessive growth factor signaling. Together, our findings suggest that HSC self-renewal is supported by cell-extrinsic mechanisms enacted by syndecan-2 from the MSC niche, highlighting the importance of the niche proteoglycome in HSC functions.

**KEY POINTS:**

- The heparan sulfate proteoglycan syndecan-2 expressed by mesenchymal stromal cells but not endothelial cells regulates HSC self-renewal
- Depletion of syndecan-2 from *Lepr*-targeted mesenchymal stromal cells remodels the hematopoietic stem cell transcriptional landscape

## INTRODUCTION

Through a careful balance of self-renewal and differentiation, hematopoietic stem cells (HSCs) maintain the hematopoietic system. HSCs reside within the bone marrow (BM) microenvironment, or niche, where they receive cues from surrounding vascular and stromal cells, which influence their functions [1-6]. BM mesenchymal and endothelial cells (ECs) produce soluble growth factors that support HSC self-renewal and differentiation at steady-state and during stress [3,4,7-14]. However, the upstream mechanisms that coordinate growth factor signals in the BM niche are incompletely understood.

Heparan sulfate proteoglycans are a class of molecules that bind growth factors to regulate their signaling potency by changing their bioavailability, creating gradients, and protecting them from degradation [15-17]. Growth factors, like stem cell factor (SCF), transforming growth factor beta 1 (TGF-β1), and stromal derived factor-1 (CXCL12), bind heparan sulfates [18-21]. We previously discovered that HSCs are highly enriched for the expression of syndecan-2 (*Sdc2*), a heparan sulfate proteoglycan, and syndecan-2 depletion reduced HSC self-renewal ability [22]. Seminal works show that heparan sulfate proteoglycans are abundantly expressed in the BM niche [23-25] and heparan sulfates regulate hematopoietic cell functions [26,27]. These findings led us to hypothesize that syndecan-2 expressed by BM niche cells regulates hematopoiesis and HSC functions.

In this study, we show that syndecan-2 is highly expressed by BM mesenchymal stromal cells (MSCs), and to a lesser degree ECs. Tissue-specific depletion of syndecan-2 from these cell compartments revealed that MSC-expressed syndecan-2 is critical for HSC self-renewal, while EC-expressed syndecan-2 is dispensable. Biochemical analyses reveal that syndecan-2 can bind hematopoietic growth factors, including TGF-β1, CXCL12, thrombopoietin (TPO), and FLT3 with nanomolar affinity. Depletion of syndecan-2 from MSCs transcriptionally reprograms HSCs to express gene signatures associated with premature exhaustion and aging, changes that are computationally predicted to occur via heparan sulfate-binding growth factor ligands. Our data extend currents models of the BM niche by revealing the critical function that the proteoglycome plays in supporting HSC self-renewal ability.

## METHODS

Detailed information regarding antibodies, instruments, reagents, and other resources are in **Supplemental Table 1**. Methods not included here can be found in Supplemental Methods.

### Mice

Animal procedures were approved by the Institutional Animal Care and Use Committee (PROTO202100049) and overseen by the Comparative Medicine Shared Resource of Fred Hutchinson Cancer Center. Mixed-sex adult mice (8-12 weeks of age) were used in the described experiments. Euthanasia was performed by CO_2_ asphyxiation followed by cervical dislocation.

### Cell isolation and analysis

For hematopoietic cells (HCs), BM was isolated from femurs and tibias by centrifugation [28] and lysed with ACK buffer. For ECs or MSCs, BM was isolated by crushing in a mortar and pestle containing liberase [11]. Isolated BM was processed in complete IMDM (IMDM: 10% FBS, 1% penicillin-streptomycin). Peripheral blood (PB) was collected via submandibular or retroorbital bleed into EDTA and subjected to complete blood count (CBC) and/or lysed with ACK buffer for cytometric analysis. BM cellularity was determined using trypan blue and an automated cell counter.

### Flow cytometry, cell sorting, and qRT-PCR

Protein expression was assessed by flow cytometry. Cells were stained live or fixed and permeabilized before analysis. Following RNA isolation, gene expression was assessed using TaqMan probes. See Supplementary Table 1 for kits, antibodies, and primers used.

### Competitive repopulation assay

Recipient SJL mice were lethally irradiated (10Gy ^137^Cs Mark 1 total body irradiator) the day before transplant. 3×10^5^ BM cells, or 250 FACS-isolated 34^-^KSL cells from *Sdc2*^ΔMSC^, *Sdc2*^ΔEC^, *Sdc2*^ΔHC^, or littermate controls (CD45.2 C57BL/6 background) were intravenously injected into recipients with 3×10^5^ (whole BM transplants) or 2×10^5^ (34^-^KSL transplants) BM cells from an SJL competitor. Donor chimerism was quantified as described [22].

### Co-culture assays

96-well U-bottom plates were pre-seeded with 4×10^3^ HS-5 human stromal cells (parent, sh*SDC2*, or shControl) per well. The next day, media was replaced with complete IMDM supplemented with TPO (20ng/ml), FLT3 (50ng/ml), and SCF (150ng/ml), and 500 34^-^KSL cells from wild-type C57BL6/J mice were added. Co-cultures were incubated for 7 days and analyzed.

### Single-cell RNA sequencing

At least 2.1×10^4^ KSL cells per mouse were FACS-isolated. Barcoding and sequencing were performed by the Fred Hutch Genomics Shared Resource.

### Statistics

Statistics were performed using Prism and R. Hypothesis tests between multiple groups or timepoints were performed using one- or two-way ANOVA followed by Holm-Sidak-corrected t-tests; tests comparing only two groups were performed using unpaired t-tests. Unless otherwise noted, data are considered significant when p<0.05.

## RESULTS

### Syndecan-2 is highly expressed by BM niche cells

Because syndecan-2 expression marks murine HSCs and supports their self-renewal potential, we interrogated syndecan family gene expression in BM niche cells. We queried published single-cell RNA sequencing (scRNA-seq) datasets examining the expression profiles of bone and BM niche cells from adult mice [29]. BM niche cells exhibit distinct expression patterns of *Sdc1, Sdc2, Sdc3*, and *Sdc4* (**Figure 1A-D**). One dataset revealed that *Sdc1* and *Sdc2* were expressed by 56% and 48% of leptin receptor^+^ MSCs, respectively, while only 6% and 41% of MSCs expressed *Sdc3* and *Sdc4* (**Figure 1B**).

**Figure 1:**
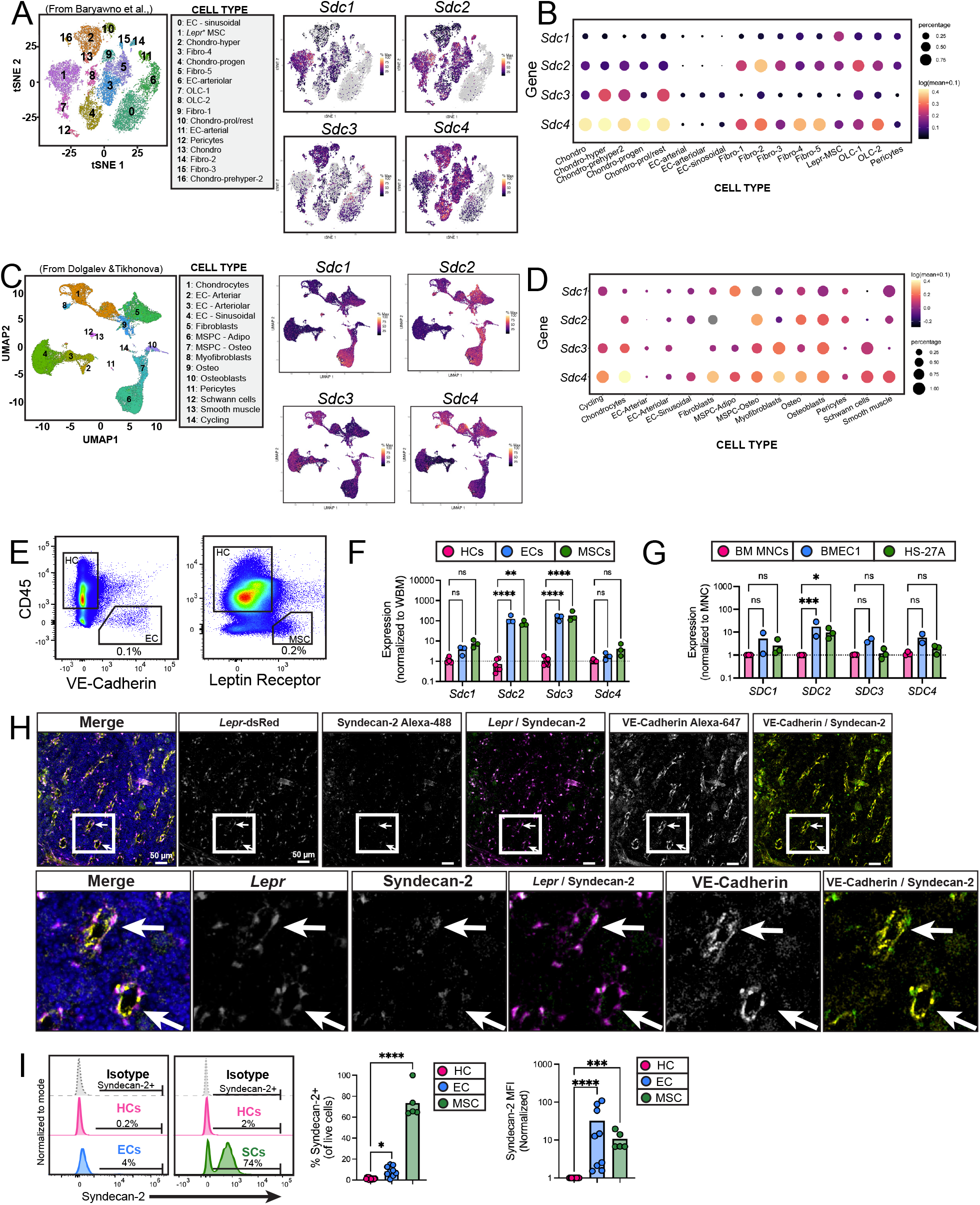
Syndecan-2 is expressed by distinct BM niche cells. **(A)** t-distributed stochastic neighbor embedding (tSNE) map of non-hematopoietic bone and bone marrow cells from data originally analyzed in PMID 31130381. Each color corresponds to a different annotated stromal cell cluster. **(B)** Expression of *Sdc* family genes within each cluster based on percent positivity and log(mean+0.1) transcripts per million (TPM). **(C)** Uniform Manifold Approximation and Projection (UMAP) visualization of bone marrow niche cells from integrated niche dataset analyses in PMID 33777933. Each color represents a different cluster, or niche cell population. **(D)** Expression of *Sdc* family genes within each cell population according to the percent positive of cells within a population and their relative expression based on log(mean+0.1) TPM. (E) Gating strategy used to isolate mouse bone marrow CD45^+^ hematopoietic cells (HCs), CD45-VE-Cadherin+ endothelial cells (ECs), and CD45^-^ Leptin Receptor^+^ mesenchymal stromal cells (MSCs). **(F)** qRT-PCR analysis of *Sdc* genes in mouse bone marrow HCs, ECs, and MSCs normalized to whole bone marrow cells (dotted line). **(G)** qRT-PCR analysis of *SDC* family genes in human bone marrow mononuclear cells, BMEC-1 endothelial cells, and HS-27A stromal cells. **(H)** Micrograph from immunofluorescence imaging of thick bone marrow sections from Lepr-cre^+^; ROSA26-tdTomato mice stained with antibodies to syndecan-2, VE-Cadherin, and DAPI. Scale bar = 50 µm. **(I)** Flow cytometric analysis of syndecan-2 surface expression on CD45^+^ HCs, CD45^-^ VE-Cadherin^+^ ECs, or CD45^-^ Leptin Receptor^+^ MSCs and quantification of % syndecan-2^+^ cells and syndecan-2 mean fluorescence intensity. (*For mouse cell qRT-PCR: n=3 independent experiments with n=3 technical replicates/experiment, statistics show Holm-Sidak’s corrected t-tests after two-way ANOVA; for human cell PCR: n=3 unique donors from n=3 independent experiments with n=3 technical replicates/experiment with averages per experiment shown, statistics show Holm-Sidak’s corrected t-tests after two-way ANOVA; for flow cytometry: n=10 independent experiments, n=5-14 biological replicates per experiment, statistics show Holm-Sidak’s corrected t-tests after one-way ANOVA. *p<0*.*05, **p<0*.*01, ***p<0*.*001, ****p<0*.*0001*.)

Other prominent *Sdc2*-expressing clusters were cell subtypes originating from bone, such as fibroblasts (fibro-4), osteolineage cells (OLC-1, OLC-2), and pericytes. Meanwhile, *Sdc2* was lowly expressed by sinusoidal ECs (sECs, 1%) and arteriolar ECs (aECs, 2%). We consulted another scRNA-seq data set that combined two independent analyses of the murine stromal BM niche composition, finding similar results (**Figure 1C-D**) [30]. *Sdc2* was strongly expressed by cells within clusters containing adipo- and osteo-lineage primed MSCs and a minority of sECs and aECs expressed *Sdc2* (**Figure 1D**). These data suggest that *Sdc2* is enriched in murine BM MSC populations relative to ECs.

We next quantified syndecan family gene expression in BM MSCs relative to BM hematopoietic cells. We used fluorescence-activated cell sorting (FACS) to isolate CD45^-^ Leptin receptor^+^ (LepR^+^) MSCs, CD45^-^ VE-Cadherin^+^ ECs, and bulk CD45^+^ HCs from the BM of adult C57BL/6 mice (**Figure 1E**). qRT-PCR analyses revealed that compared to HCs, both MSCs and ECs expressed significantly higher *Sdc2* and *Sdc3*, but not *Sdc1* and *Sdc4* (**Figure 1E-F**). To determine whether this enrichment is conserved in human niche cells, we analyzed syndecan family gene expression in primary human BM mononuclear cells relative to the BMEC-1 BM microvascular EC cell line and HS-27A BM stromal cell line. Indeed, *SDC2* was significantly enriched in both BMEC-1 cells and HS-27A cells compared to BM mononuclear cells; enrichment was not detected in *SDC1, SDC3*, or *SDC4* (**Figure 1G**). Together, these findings indicate that *SDC2* is transcriptionally enriched in murine and human BM MSCs and ECs compared to HCs.

Because gene expression does not always align with protein, we used immunohistochemistry to visualize syndecan-2-expressing cells within the murine BM relative to *Lepr*^+^ cells and VE-Cadherin^+^ ECs. Immunohistochemical staining and confocal imaging of thick femur sections from anti-VE-Cadherin infused adult *Lepr-Cre*^*+*^; *ROSA26-TdTomato* mice revealed syndecan-2 expression throughout the BM niche (**Figure 1H**) [9,31]. Magnified images depicting VE-Cadherin^+^ ECs showed syndecan-2 (green) accumulation near VE-Cadherin^+^ ECs (yellow) without obvious colocalization (**Figure 1H, bottom panel**). Instead, syndecan-2 overlays strongly with the *Lepr*-expressing cells (magenta), as is apparent by the white color combination. Therefore, the BM expression of syndecan-2 appears to be primarily perivascular and confined to locations near *Lepr*-expressing cells. To confirm these findings, flow cytometric analysis of BM cells showed that 1.3 ± 0.9% of bulk HCs and 8.4 ± 4.8% sECs express syndecan-2 on the cell surface (**Figure 1I**). Meanwhile, MSCs exhibited a nearly tenfold increased fraction of cells expressing syndecan-2 (73.5 ± 15.3%) compared to ECs and a more than 100-fold increase compared to HCs. Mean fluorescence intensity analyses indicate significantly enriched syndecan-2 expression on ECs and MSCs compared to HCs (**Figure 1I**). Together, these findings reveal that syndecan-2 is expressed by BM ECs and MSCs, the function of which is unknown.

### Steady-state hematopoietic balance is regulated by cell-specific syndecan-2 expression

To test the role of syndecan-2 expressed by HCs, ECs, and MSCs in hematopoiesis, we used a *Sdc2*^*flox/flox*^ mouse strain. *Sdc2* was conditionally depleted in *Vav1*-targeted cells (HCs), *Cdh5*-targeted cells (ECs), and *Lepr*-targeted cells (MSCs) (**Supplemental Figure 1A**). Mice were produced by breeding hemizygotes, and we used litter-matched promoter-specific Cre negative (Cre-) as Control mice for each strain throughout our studies. This approach generated *Vav1*-cre^+^; *Sdc2*^*flox/flox*^ (*Sdc2*^ΔHC^), *Cdh5*-cre^+^; *Sdc2*^*flox/flox*^ (*Sdc2*^ΔEC^), and *Lepr*-cre^+^; *Sdc2*^*flox/flox*^ mice (*Sdc2*^ΔMSC^) mice, which had significantly lower syndecan-2 surface expression compared to Control mice (**Supplemental Figure 1B-D**).

CBC analyses of the PB of adult mice showed that *Sdc2*^Δ*MSC*^ mice had significantly increased PB white blood cells (WBCs) and lymphocytes compared to Control mice, while neutrophils, monocytes, eosinophils, and basophils were not significantly altered (**Figure 2A**). Red blood cells (RBCs), platelets, and hemoglobin levels were also unchanged in *Sdc2*^ΔMSC^ mice compared to Control mice (**Figure 2B**). Flow cytometric analysis of the PB revealed significantly elevated frequencies and numbers of B cells in *Sdc2*^Δ*MSC*^ compared to Control mice, while T and myeloid cells were unchanged (**Figure 2C-E**). These findings suggest that MSC-derived syndecan-2 contributes to PB hematopoietic balance in the B cell compartment.

**Figure 2:**
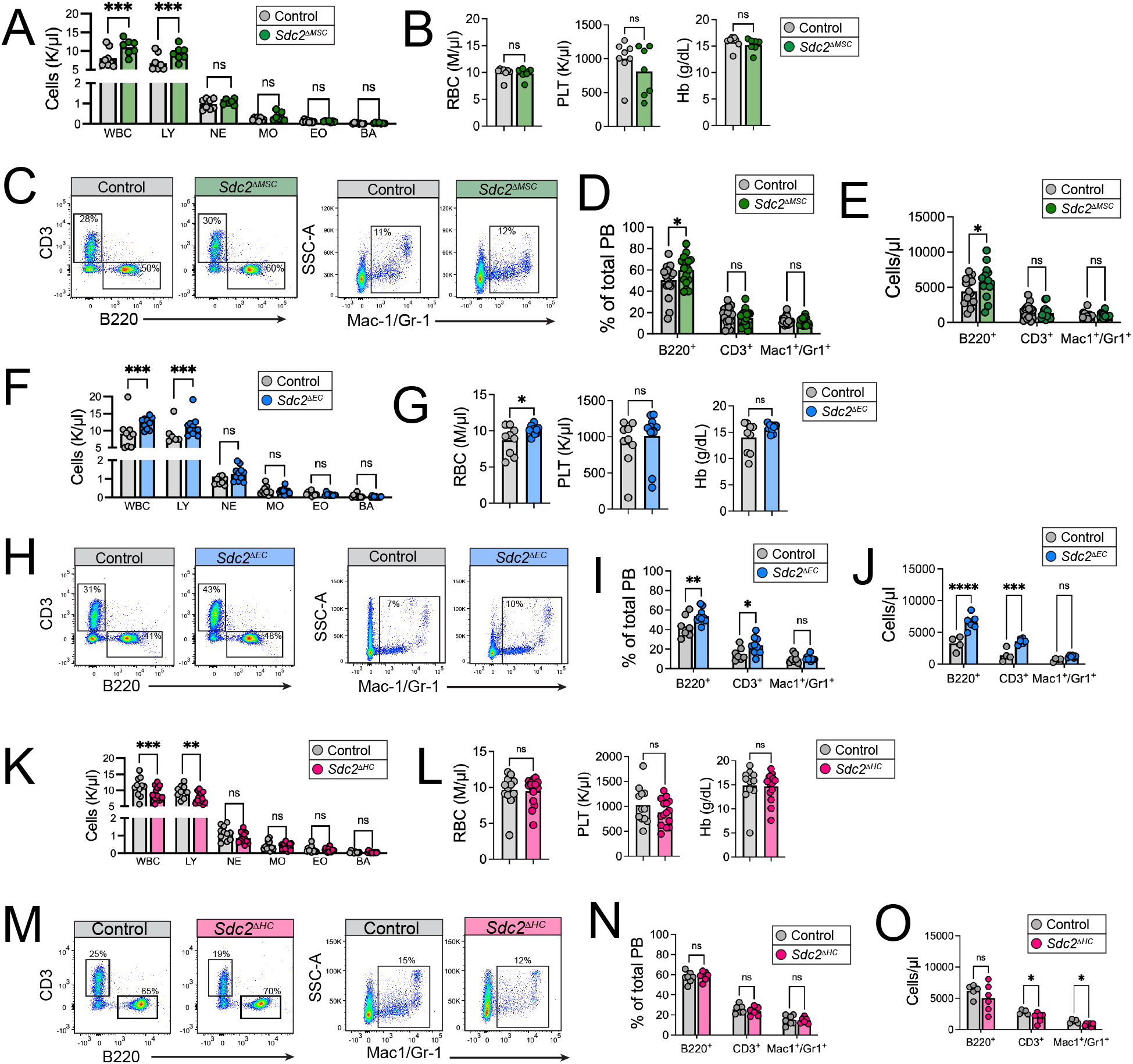
Decreased hematopoietic cell-intrinsic and extrinsic syndecan-2 skews hematopoietic balance. Complete blood count analysis of peripheral blood from 8-12-week-old mixed sex Control or *Sdc2*^ΔMSC^ mice for **(A)** white blood cells (WBC), lymphocytes (LY), neutrophils (NE), monocytes (MO), eosinophils (EO), and basophils (BA) and **(B)** red blood cells, platelets (PLT) and hemoglobin (Hb). **(C)** Representative flow cytometry gating of peripheral blood CD3^+^ T cells, B220^+^ B cells, and Mac-1/Gr-1^+^ myeloid cells and quantification of the **(D)** frequency and **(E)** number of cells in the peripheral blood. **(F)** Complete blood count analysis of peripheral blood from 8-12-week-old mixed sex Control or *Sdc2*^ΔEC^ mice for white blood cell populations and **(G)** RBCs, PLT, and Hb content. **(H)** Representative flow cytometry scatter plots showing gating strategy for peripheral blood CD3^+^ T cells, B220^+^ B cells, and Mac-1/Gr-1^+^ myeloid cells and quantification of cell **(I)** frequency and **(J)** number in the peripheral blood. Complete blood count analysis of peripheral blood from 8-12-week-old mixed sex Control or *Sdc2*^ΔHC^ mice for **(K)** white blood cells and **(L)** RBC, PLT, and Hb levels. **(M)** Representative flow cytometry gating used to identify CD3^+^ T cells, B220^+^ B cells, and Mac-1/Gr-1^+^ myeloid cells, and cell **(N)** frequencies and **(O)** numbers in the peripheral blood. (*For CBCs: n=7-15 biological replicates/strain, statistics show Holm-Sidak corrected t-tests after two-way ANOVA or for single comparisons, unpaired t-tests; for PB analyses: n=5-16 biological replicates/strain; statistics show Holm-Sidak corrected t-tests after two-way ANOVA. *p<0*.*05, **p<0*.*01, ***p<0*.*001*.)

*Sdc2*^Δ*EC*^ mice also had significantly increased PB WBCs and lymphocytes compared to Control mice, while neutrophils, monocytes, eosinophils, and basophil levels were unaltered (**Figure 2F**). *Sdc2*^Δ*EC*^ had elevated RBCs, but unchanged platelets and hemoglobin levels compared to Control mice (**Figure 2G**). PB flow cytometric analysis demonstrated that B and T cell frequencies and numbers were increased in *Sdc2*^Δ*EC*^ mice compared to Controls, while myeloid cells were unchanged (**Figure 2H-J**). Together, these findings show *Sdc2* depletion in *Lepr*-targeted and *Cdh5*-targeted cells similarly impact peripheral hematopoietic balance in the lymphoid compartment, but their contributions to specific lymphoid lineages varies.

While only ∼2% of bulk HCs express syndecan-2 (**Figure 1I**), *Sdc2*^Δ*HC*^ mice exhibited significantly decreased PB WBC, and lymphocytes compared to Control mice (**Figure 2K**). Meanwhile, neutrophils, RBC, platelets, and hemoglobin were unchanged (**Figure 2L**). PB flow cytometric lineage analyses did not detect lineage skewing but revealed significantly fewer CD3^+^ T cells and Mac-1/Gr-1^+^ myeloid cells compared to Controls (**Figure 2M-O**). These findings suggest that HC syndecan-2 supports PB hematopoietic cellularity and this contribution opposes that of EC and MSC syndecan-2.

Due to the interplay between BM and PB in regulating hematopoiesis, we analyzed the BM of our transgenic mice. Gross histological BM structure and cellularity were unchanged upon depletion of *Sdc2* from MSCs, ECs, or HCs (**Supplemental Figure 2A-F)**. BM lineage cell frequencies were also unchanged in Control mice compared to *Sdc2*^ΔMSC^, *Sdc2*^ΔEC^ or *Sdc2*^ΔHC^ mice (**Supplemental Figure 2G-I**). We detected no difference in the frequency of common myeloid, granulocyte/macrophage, megakaryocyte/erythroid, or common lymphoid progenitor cells upon *Sdc2* depletion in MSCs, ECs, or HCs (**Supplemental Figure 2J-L**). The percent and abundance of LT-HSCs were also unchanged (**Supplemental Figure 2M-O**). Therefore, syndecan-2 from MSCs, ECs, and HCs does not appear to regulate BM hematopoietic cell balance or numbers.

### Short-term hematopoietic repopulation is unaffected by syndecan-2 depletion from MSCs, ECs, or HCs

As our prior works indicate that HSC-intrinsic syndecan-2 supports HSC repopulation ability and self-renewal, we sought to test whether syndecan-2 from MSCs, ECs, or bulk HCs has a similar function. To test for cell-source-specific HSC functional defects, we performed competitive repopulation assays. We transplanted whole BM (WBM) cells from *Sdc2*^Δ*MSC*^ *and* Control mice (CD45.2) against WBM cells from SJL mice (CD45.1) into lethally irradiated SJL recipient mice (**Figure 3A**). Transplant of Control or *Sdc2*^ΔMSC^ cells yielded similar levels of PB donor chimerism overall, or within B, T, or myeloid, compartments (**Figure 3B**) and donor chimerism in the BM after 16 weeks (**Supplemental Figure 3A**). Similar results were obtained using donor cells from *Sdc2*^ΔEC^ and *Sdc2*^ΔHC^ strains for competitive transplants, whereby PB donor chimerism (**Figure 3C-F**) and BM donor chimerism (**Supplemental Figure 3B-C**) did not differ significantly from litter-matched control mice. Together, these data suggest that syndecan-2 from MSCs, ECs, or HCs do not contribute to short-term HSC repopulation ability, in agreement with our previous findings that HSC-derived syndecan-2 does not control ST-HSC repopulation capacity [22].

**Figure 3:**
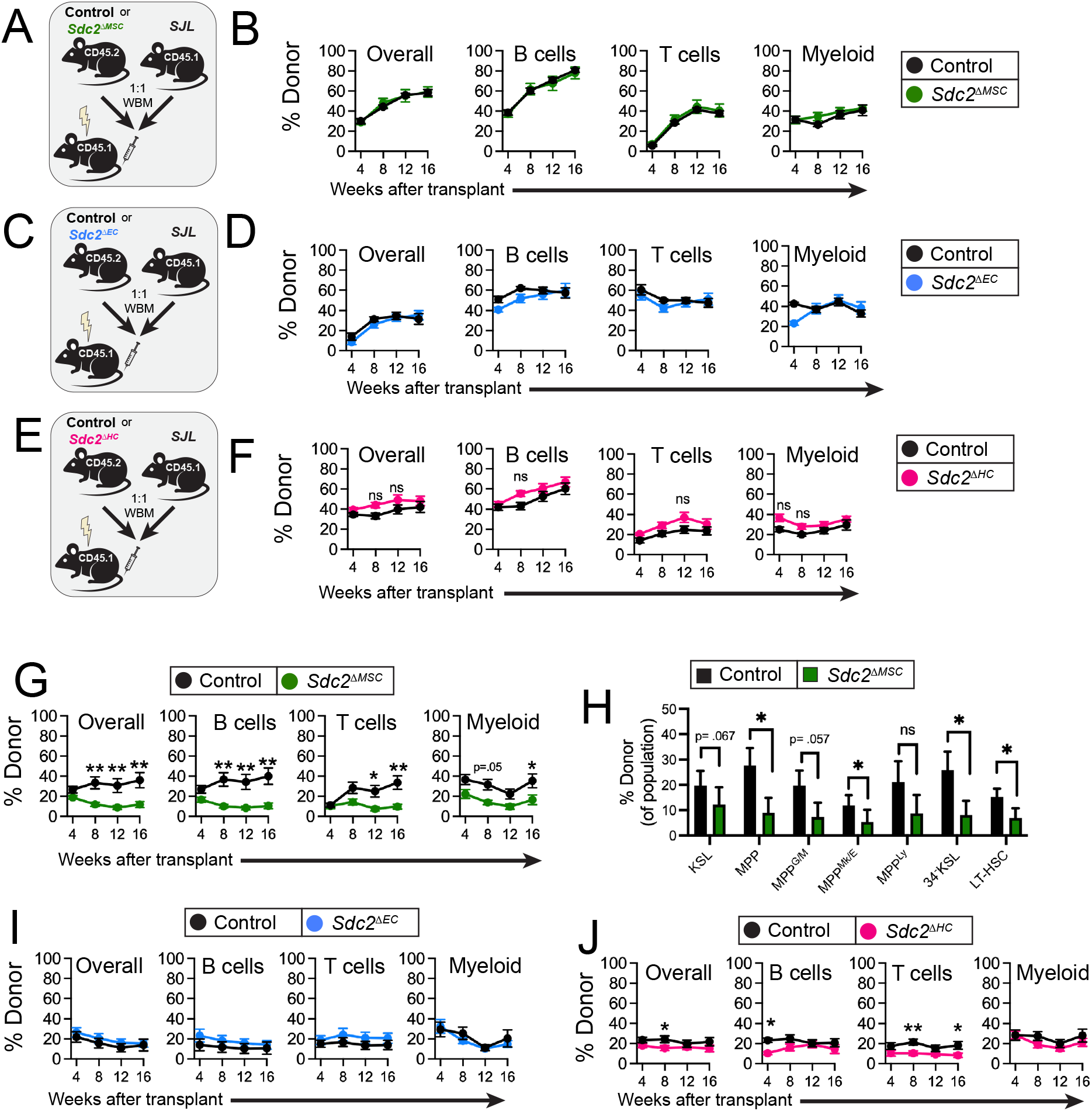
Depletion of *Sdc2* from *Lepr*-targeted cells, but not *Cdh5*-targeted cells reduces HSC self-renewal ability. **(A)** Experimental design for primary competitive repopulation assay using bone marrow cells from Control or *Sdc2*^ΔMSC^ mice as donors. **(B)** Peripheral blood CD45.2 donor cell chimerism overall and within the B220+ B cell, CD3+ T cell, and Mac-1/Gr-1 myeloid cell compartments after primary competitive transplant. **(C)** Experimental design for primary competitive repopulation assay using Control or *Sdc2*^ΔEC^ mice as donors. **(D)** Peripheral blood CD45.2 donor cell chimerism overall and within the B cell, T cell and myeloid cell compartments after primary transplant. **(E)** Experimental design for primary competitive repopulation assay using Control or *Sdc2*^ΔHC^ mice as donors. **(F)** Peripheral blood CD45.2 donor cell chimerism overall and within the B cell, T cell, and myeloid cell compartments after primary transplant. **(G)** Overall and multilineage peripheral blood donor cell chimerism after secondary transplant of whole bone marrow cells from Control or *Sdc2*^ΔMSC^ mice and **(H)** bone marrow donor hematopoietic stem/progenitor cell chimerism. **(I)** Overall and multilineage peripheral blood donor cell chimerism after secondary transplant of whole bone marrow cells from Control or *Sdc2*^ΔEC^ or **(J)** *Sdc2*^ΔHC^ mice. (*For primary transplants, n=10 recipients/strain; for secondary transplants n=7-10 recipients/strain; statistics show Holm-Sidak corrected t-tests after two-way ANOVA. *p<0*.*05, **p<0*.*01*.).

### Syndecan-2 from MSCs, but not ECs, supports long-term HSC self-renewal

We next performed secondary competitive transplantations, using WBM from recipient mice that received primary transplantation of Control or *Sdc2*^Δ*MSC*^ WBM donor cells. We detected significantly diminished overall donor chimerism upon transplant of *Sdc2*^Δ*MSC*^ cells compared to Control cells, concomitant with decreased B, T, and myeloid donor cell contribution (**Figure 3G**). BM analyses revealed significantly decreased overall and lineage (**Supplemental Figure 3D**) and LT-HSC and 34^-^KSL (**Figure 3H**) donor chimerism in *Sdc2*^Δ*MSC*^ donors compared to Control mice. There was also significantly reduced donor chimerism within the multipotent progenitor (MPP) and MPP megakaryocyte/erythroid (MPP^Mk/E^) populations, indicating that HSCs from MSC environments deficient in syndecan-2 have decreased self-renewal ability and progenitor maintenance capacity. To test whether this effect of syndecan-2 on HSC self-renewal extends to EC- and HC-derived syndecan-2, we performed analogous secondary transplant studies using *Sdc2*^ΔEC^ or *Sdc2*^ΔHC^ mice. PB and BM analyses revealed that *Sdc2*^Δ*EC*^ and Control mice had similar donor cell chimerism upon secondary transplantation (**Figure 3I, Supplemental Figure 3E**), suggesting EC syndecan-2 is dispensable for HSC self-renewal. However, *Sdc2*^Δ*HC*^ mice had decreased overall, B, and T PB donor cell chimerism compared to Control mice (**Figure 3J**), with BM chimerism unchanged (**Supplemental Figure 3F**). The effect size and persistence throughout the transplant of *Sdc2*^ΔHC^ cells, however, were much more modest than the depletion of MSC-derived syndecan-2, suggesting that the contribution of syndecan-2 from the MSC niche to HSC self-renewal may be more substantial than that of HCs.

### HSCs from mice depleted of syndecan-2 from MSCs exhaust more rapidly upon transplant

To better pinpoint the effects of MSC-derived syndecan-2 on HSCs, we performed competitive transplants using FACS-sorted 34^-^KSL cells from *Sdc2*^ΔMSC^ mice or Control mice as donor cells (**Figure 4A**). This approach enables us to examine the effects of HSCs in the absence of other donor hematopoietic progenitor cells or lineage cells, which could influence HSCs upon transplant. PB donor chimerism was unchanged overall in primary transplants, but *Sdc2*^ΔMSC^ HSCs yielded significantly reduced donor myeloid cell chimerism compared to Controls (**Figure 4B**), in agreement with our prior work showing *Sdc2* knockdown in HSCs reduces myeloid repopulation [22]. Further, overall BM donor chimerism was significantly reduced in mice transplanted with 34^-^KSL cells from *Sdc2*^ΔMSC^ mice compared to Controls, accompanied by decreased BM myeloid chimerism, while B and T cell chimerism was unchanged (**Figure 4C**). Donor-derived LT-HSCs and progenitor cells in the BM of mice transplanted with 34^-^KSL cells from *Sdc2*^ΔMSC^ mice were significantly lower compared to Control mice (**Figure 4D**). These data suggest that HSCs from *Sdc2*^ΔMSC^ mice exhaust more rapidly upon competitive transplant. Consistent with these findings, secondary transplants revealed significantly decreased PB and BM multilineage donor cell chimerism (**Figure 4E-F**). Together, these data provide further evidence that that syndecan-2 from MSCs supports HSC self-renewal ability.

**Figure 4:**
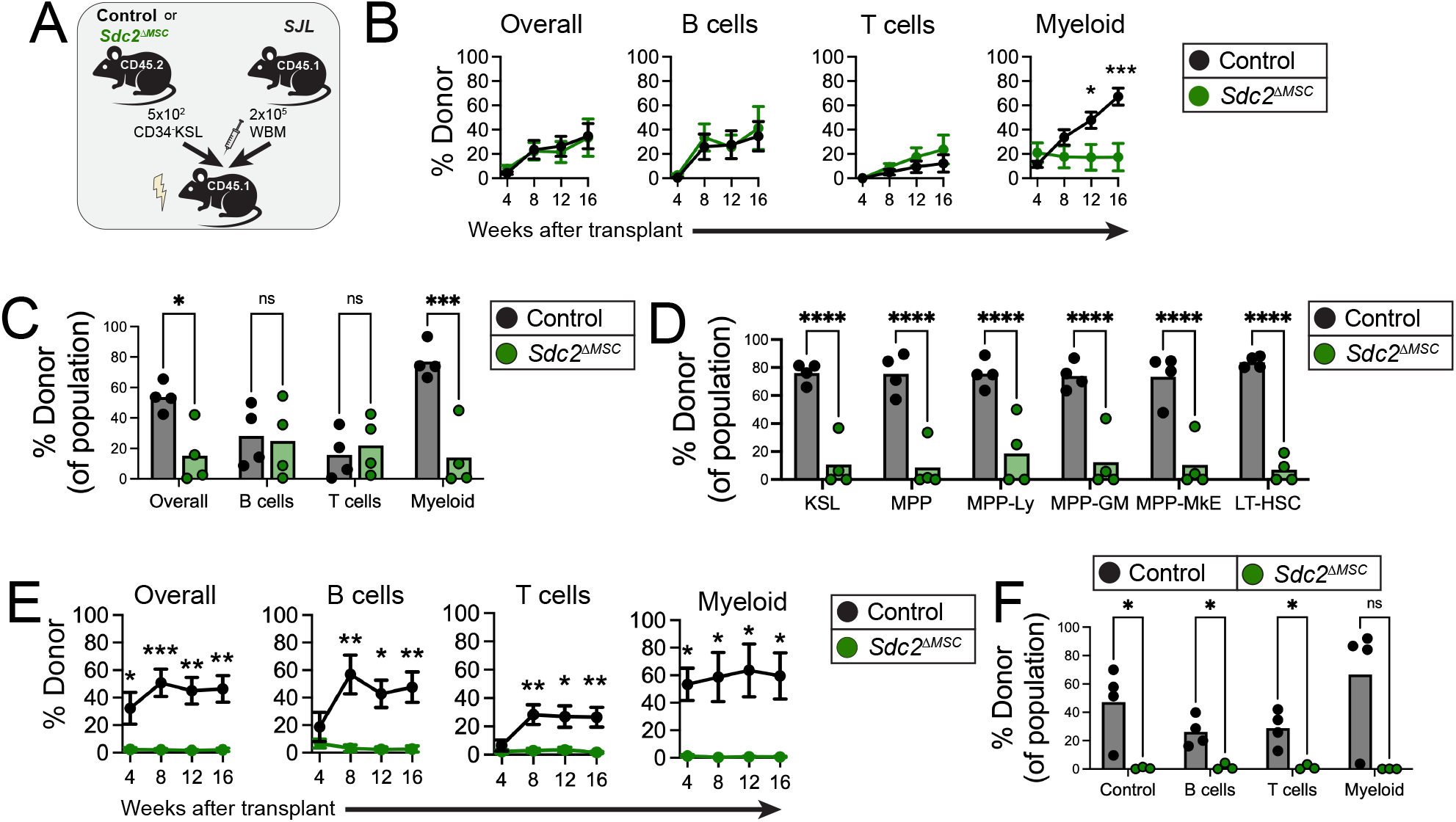
HSCs from mice depleted of syndecan-2 from MSCs exhaust more rapidly upon transplant. **(A)** Experimental design for primary competitive repopulation assay using 34^-^KSL cells from Control or *Sdc2*^ΔMSC^ mice as donor cells. **(B)** Peripheral blood and **(C)** bone marrow CD45.2 donor cell chimerism overall and within the B220^+^ B cell, CD3^+^ T cell, and Mac-1/Gr-1^+^ myeloid cell compartments after primary competitive transplant. **(D)** Quantification of bone marrow hematopoietic stem/progenitor cell chimerism at 16-weeks post-transplant. **(E)** Overall and multilineage peripheral blood CD45.2 donor cell chimerism after secondary transplant of donor-derived whole bone marrow cells from primary transplanted recipient mice in (A) and **(F)** bone marrow donor cell chimerism at 16-weeks post-secondary transplant. (*n=4 recipients/strain for primary and secondary transplants; statistics show Holm-Sidak corrected t-tests after two-way ANOVA. *p<0*.*05, **p<0*.*01, ***p<0*.*001, ****p<0*.*0001*.).

To further explore the role of MSC-derived *Sdc2* on HSCs, we tested the impact on wild-type hematopoietic cells. We transplanted a radioprotective dose of whole BM cells into lethally irradiated Control or *Sdc2*^ΔMSC^ mice (**Supplemental Figure 4A**). We detected no change in CBCs, PB lineage cells, or BM lineage cells at 24-weeks post-transplant (**Supplemental Figure 4B-D**). Further, we did not identify changes in BM MPP, MPP^G/M^, MPP^Mk/E^, or LT-HSC numbers upon hematopoietic transplant into Control or *Sdc2*^ΔMSC^ mice (**Supplemental Figure 4E**). These data suggest that adult wild-type HSCs may be able to withstand the influence of the *Sdc2*^ΔMSC^ niche over extended time periods. These data suggest that the impact of the *Sdc2*^ΔMSC^ niche on HSC functionality may operate in a temporally restricted manner.

### Loss of syndecan-2 from MSCs promotes HSC exhaustion

Because our studies generating chimeric mice in Supplemental Figure 4A required Control and *Sdc2*^ΔMSC^ mice to be irradiated before transplant, and irradiation is known to substantially remodel the BM niche, we sought to use a more simplified approach to test how syndecan-2 from the stroma influences HSCs. To analyze whether syndecan-2 from the stroma directly influences HSCs, we co-cultured HSCs with the HS-5 BM-derived stromal cell line for 7 days [32]. We transduced HS-5 cells with lentiviral particles containing Control or *SDC2* targeting shRNAs to generate shControl or sh*SDC2* cells, which have significantly reduced *SDC2* expression compared to shControl cells (**Figure 5A**). *Ex vivo* HSCs culture in cytokine-enriched media is a stress model that stimulates HSC proliferation and exhaustion, enabling us to rapidly screen for stromal-mediated effects of *SDC2* on HSC maintenance [33-36]. We therefore cultured 34^-^KSL cells from C57BL/6 mice with Control or sh*SDC2* HS-5 cells in growth factor-enriched expansion media containing TPO, SCF, and FLT3 ligand (TSF media) (**Figure 5B**). After 7 days of culturing, we quantified hematopoietic stem/progenitor cells using flow cytometry. Culturing 34^-^KSL cells with sh*SDC2* cells resulted in a significantly greater frequency of Lin^+^ progeny concomitant with decreased Lin^-^ cells, KSL cells, and CD34^-^KSL cells compared to co-culture with shControl cells (**Figure 5C-D**). 34^-^KSL expansion relative to input cells was also significantly reduced upon co-culture with sh*SDC2* cells compared to shControl cells (**Figure 5E**). Together, these data indicate that depletion of stromal syndecan-2 reduces HSC expansion ability and causes HSCs to exhaust more rapidly.

**Figure 5:**
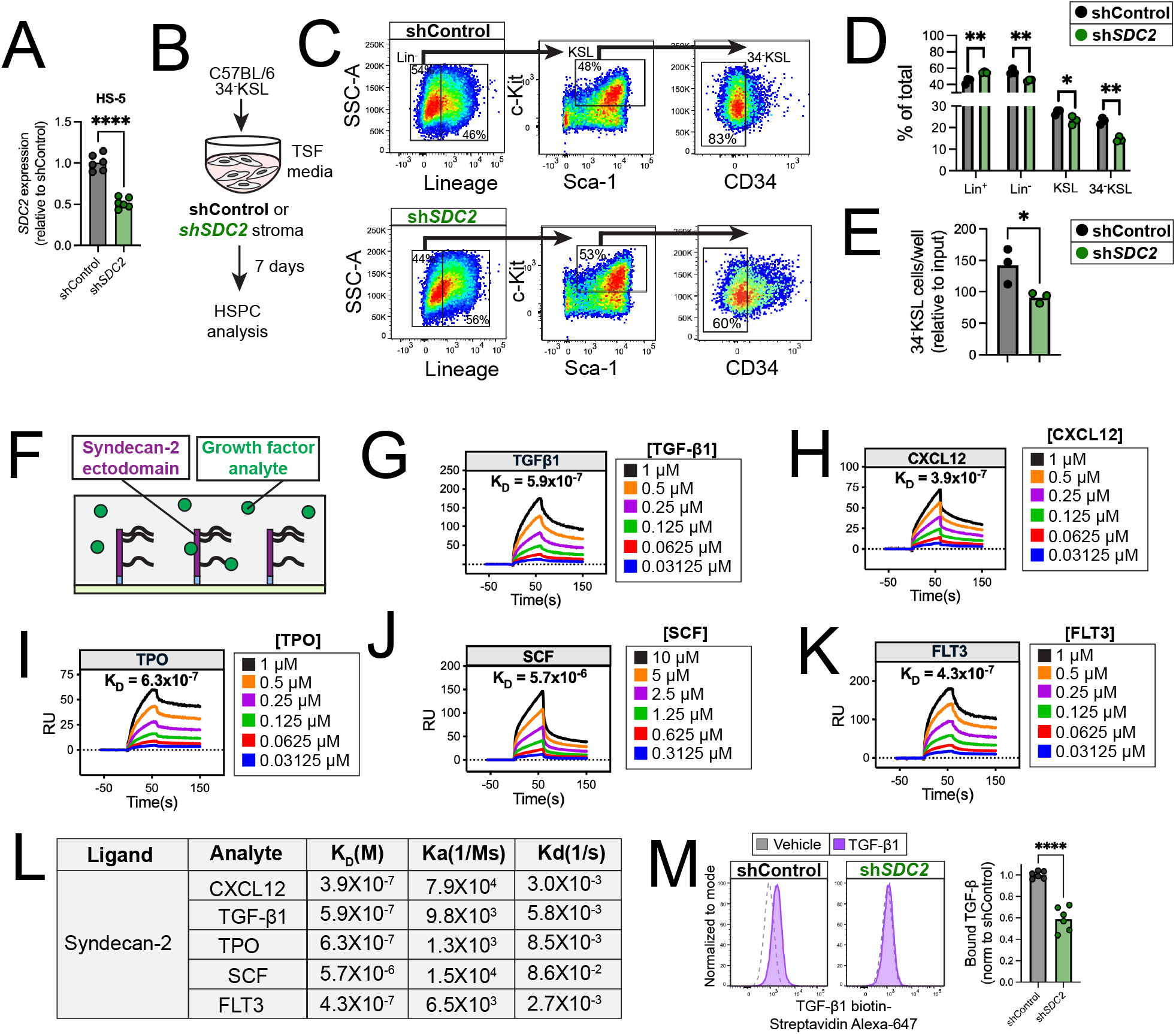
Syndecan-2 binds growth factors and facilitates hematopoietic responses to growth factor stimuli. **(A)** qRT-PCR quantification of *SDC2* expression in shControl and sh*SDC2* HS-5 cells. **(B)** Experimental design used for co-culturing 34^-^KSL cells with HS-5 cells in TSF media. **(C)** Representative scatter plots showing gating strategy used to analyze hematopoietic cell populations at day +7 after co-culture. **(D)** Quantification of the frequency of Lin^+^, Lin^-^, KSL, and 34^-^KSL cells after co-culture and **(E)** the number of 34^-^KSL cells per well. **(F)** Experimental design for Surface Plasmon Resonance analysis of protein interactions. **(G-K)** Sensograms depicting binding kinetics of syndecan-2 ectodomain to **(G)** TGF-β1, **(H)** CXCL12, **(I)** TPO, **(J)** SCF, and **(K)** FLT3 and **(L)** binding properties. **(M)** Representative histograms quantifying TGF-β1 bound to shControl and sh*SDC2* HS-5 cells from flow cytometric analyses. (*For co-culture studies: n=3 replicates/cell line; statistics show unpaired t-tests, error bars show SEM; for binding assays: n=3 independent experiments with n=2 technical replicates per experiment, statistics show unpaired t-test. *p<0*.*05, ****p<0*.*0001*.).

### Syndecan-2 binds hematopoietic growth factors

Hematopoietic growth factors, like SCF, TGF-β, and CXCL12 are known to bind heparan sulfate glycan chains. We reasoned that syndecan-2 from MSCs may bind growth factors to change their signaling potency for HSCs. We first sought to determine whether syndecan-2 interacts with these growth factors, and if so, define the properties of such interactions. We used surface plasmon resonance (SPR) to quantify purified murine syndecan-2 ectodomain binding to TGF-β1 or CXCL12, or growth factors enriched in TSF media: TPO, SCF, or FLT3 (**Figure 5F**). SPR sensograms show that TGF-β1, CXCL12 TPO, SCF, and FLT3 associate with syndecan-2, and the level of association increases with higher ligand concentration (**Figure 5G-K**). We used these data to calculate the dissociation constant, K_D_, finding that syndecan-2 has nanomolar binding affinity to CXCL12 and TGF-β1, and micromolar affinity to SCF (**Figure 5L**). Quantification of the association rate constant, K_a_, indicates that syndecan-2 binds SCF more slowly than CXCL12 and TGF-β. Further, the dissociation rate constant, K_d_, is also higher for SCF than CXCl12 and TGF-β1, suggesting that syndecan-2 has more dynamic interactions with SCF than CXCL12 and TGF-β1.

To build on these molecular interaction data, we tested whether syndecan-2 regulates growth factor binding to the HS-5 BM-derived stromal cell line. We treated HS-5 cells with biotinylated TGF-β1 and quantified binding with streptavidin detection using flow cytometry. TGF-β1 binding to HS-5 cells depleted of *SDC2* was significantly lower than to Control cells (**Figure 5M**). These data indicate that syndecan-2 regulates the ability for growth factors to bind stromal cells.

### Deficiency of syndecan-2 from MSCs remodels the HSC transcriptome

We next used scRNA-seq to understand how syndecan-2 from MSCs regulates HSCs at the transcriptional level. We isolated KSL hematopoietic stem and progenitor cells from the BM of adult Control or *Sdc2*^ΔMSC^ mice for scRNA-seq analysis (**Figure 6A**). Clustering and Garnett-guided, supervised cell type annotation revealed nine hematopoietic cell groups, including an HSC group (**Figure 6B**). There were 82 differentially expressed genes in HSCs from *Sdc2*^ΔMSC^ mice compared to HSCs from Control mice, including *Fosb, Fos, Jun, Jund, Dusp1, Erg1, Rhob*, and *Il31ra* (**Figure 6C**). Metascape analysis revealed significant enrichment in gene sets associated with cellular response to growth factor stimulus and cytokine-mediated signaling pathways in *Sdc2*^ΔMSC^ HSCs compared to Control HSCs (**Figure 6D**) as well as enrichment in gene sets associated with mononuclear cell and leukocyte differentiation. Because many of these differentially expressed genes are associated with aging, we analyzed HSC expression of a signature recently defined to be enriched in old HSCs [37]. *Sdc2*^ΔMSC^ HSCs were significantly enriched for the gene signature associated with old HSCs compared to Control HSCs (**Figure 6E**). These data show that the transcriptional profile of HSCs cultivated in niches depleted of *Sdc2* from *Lepr*-targeted cells are significantly different from Control HSCs.

**Figure 6:**
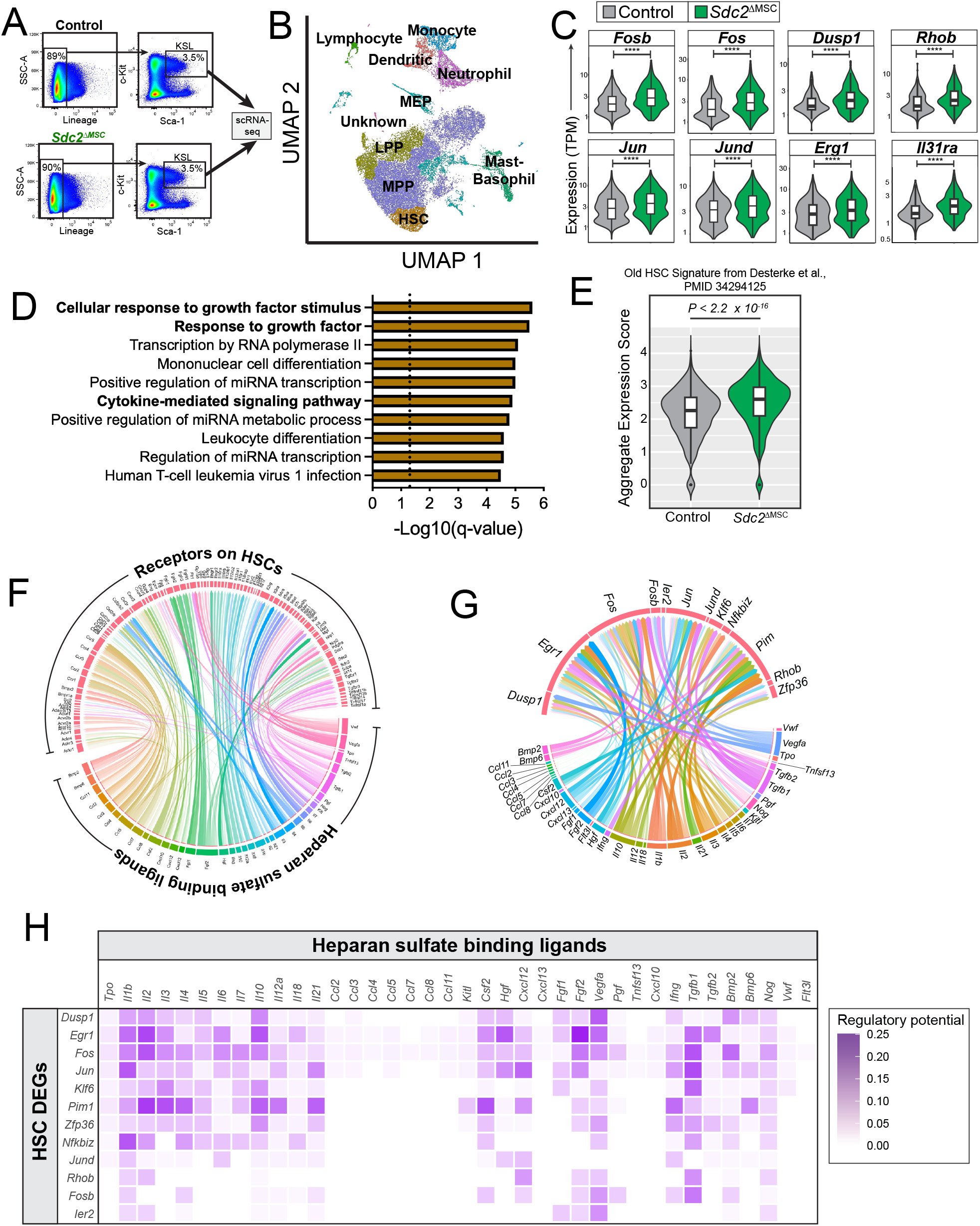
Stromal syndecan-2 remodels long-term hematopoietic stem cell transcriptome. **(A)** Gating strategy used to isolate bone marrow 34^-^KSL cells from Control or *Sdc2*^ΔMSC^ mice, which we subjected to single-cell RNA sequencing. **(B)** UMAP visualization of isolated cells and post-hoc classification; each color represents a different population. **(C)** Selected expression profiles of differentially expressed genes (DEGs) in HSCs from Control and *Sdc2*^ΔMSC^ mice. **(D)** Metascape top terms for DEGs identified using a linear regression fit to a quassipoisson distribution (Monocle 3 default). **(E)** Box plot of old HSC signature expression of all HSCs from the Control and *Sdc2*^ΔMSC^ mice (n=2 mice per genotype). The significance of differences in the two conditions was calculated using a Wilcoxon rank sum test. **(F)** Circos plot depicting heparan sulfate-binding ligand – HSC receptor interactome. **(G)** Circos plot showing interactions between heparan sulfate-binding ligands and DEGs (salmon); transparency is proportional to regulatory potential. **(H)** NicheNet ligand –target matrix showing the regulatory potential between heparan sulfate-binding ligands and DEGs differentially expressed in *Sdc2*^ΔMSC^ HSCs compared to Control HSCs. (*All differentially expressed genes have a q-value (adjusted p-value) < 0*.*05; regulatory potential is averaged for HSCs from n=2 mice/strain sequenced in duplicate*)

The defined function of heparan sulfate proteoglycans, like syndecan-2 is to bind growth factors and regulate their signaling potency. Therefore, we used the NicheNet algorithm to quantify the regulatory potential of specific heparan sulfate-binding ligand-receptor interactions for controlling the abundance of differentially expressed genes in *Sdc2*^ΔMSC^ HSCs compared to Control HSCs [38,39]. Heparan sulfate-binding ligand-receptor pairs had varying levels of regulatory potential for the differentially expressed genes we detected in the HSC cluster (**Figure 6F**). Further evaluation revealed that *Tgfb1, Fgf2*, and *Il2* were among the top ligands with regulatory potential for *Fos* expression, while *Cxcl12* and *Il1b* had strongest regulatory potential for *Jun* (**Figure 6G-H**). Taken together, these findings suggest that *Sdc2* from *Lepr*-targeted cells regulates HSC transcriptional signatures, which is predicted to occur via heparan-sulfate-binding growth factors we determined to directly bind syndecan-2.

## DISCUSSION

Our results reveal that syndecan-2 depletion causes hematopoietic system imbalances and HSC defects in a cell source specific manner. Intriguingly, our work shows that syndecan-2 is highly expressed by BM MSCs, but lower in BM ECs. While there is no change in HSC number in HSCs freshly isolated from niches depleted of syndecan-2 in MSCs, they exhaust more rapidly than Control HSCs in competitive transplants. These data imply that at steady-state, the adult hematopoietic system can withstand the impact of syndecan-2 depletion from MSCs. However, in the context of a stress, like a hematopoietic cell transplant or *ex vivo* culture, HSCs have incurred deficiencies that cannot be corrected even with support provided by cytokines or a wild-type environment. Future studies evaluating whether this holds true in the context of hematopoietic aging, a system with HSC fitness declining naturally, will enable us to understand the limitations of such HSC adaptations.

HSCs highly express syndecan-2 on their cell surface, which led us to theorize that the expression of syndecan-2 in the microenvironment may be obsolete for HSCs, as they have their own intrinsic regulatory mechanisms in place via syndecan-2. However, our data also indicates that despite expressing syndecan-2 themselves, HSCs still depend on syndecan-2 from their MSC niche for their self-renewal ability. Whether these HSC-intrinsic and HSC-extrinsic functions of syndecan-2 contribute to hematopoiesis and HSC self-renewal throughout the lifetime of an organism requires further studies.

At a transcriptional scale, HSCs respond to depletion of syndecan-2 in the MSC compartment by increasing their expression of *Fosb, Fos, Jun*, and *Jund*. Fos and jun compose the activator protein 1 (AP-1) transcription factor; their increased expression is associated with impaired HSC repopulating ability [40]. However, the role of *Fosb* in HSC functions is context dependent. Prolonged expression of *c-fos* can reduce BM Lin^-^Sca-1^+^ cell cycle entry and differentiation ability [41], while c-jun downregulation increased engraftable HSC numbers upon *ex vivo* expansion [42]. Meanwhile, HSCs cultured in hibernation conditions depleted of SCF and TPO express lower *Fosb* than freshly isolated HSCs [43]. Further, However, *Slc23a2-*deficient HSCs, which have increased self-renewal ability compared to their control counterparts, express increased *Fosb* [44]. Therefore, while the expression of AP-1 transcription factors in HSCs is context-dependent, conditions that disrupt these pathways significantly impact HSC maintenance and self-renewal.

Heparan sulfate-binding growth factors like SCF and TGF-β can promote AP-1 activation [45,46], consistent with findings from our scRNA-seq studies that HSCs from *Sdc2*^ΔMSC^ mice were significantly enriched for gene sets associated with growth factor and cytokine-mediated signaling. However, future mechanistic studies are needed to determine whether syndecan-2 tempers HSC responses to these growth factors directly or indirectly. By binding growth factors, heparan sulfate proteoglycans create a reservoir of biologically active molecules, which can protect cells from hyperstimulation. Our data indicate that syndecan-2 binds several growth factors critical for HSCs, including SCF, TPO, CXCL12, and TGF-β [4,13,47-50]. Therefore, the effects of syndecan-2 on hematopoiesis may be the result of controlling the localization and potency of several growth factors. Further, the predominant function of syndecan-2 in the BM niche may depend on the local concentration of heparan sulfate binding molecules. Saturation of syndecan-2-binding sites may render syndecan-2 unable to organize the BM growth factor milieu, but further investigation is needed to support this concept.

In summary, we discovered that syndecan-2 expressed by MSCs regulates HSC self-renewal ability, responses to growth factor stimuli, and transcriptional signatures. Future work is needed to determine how we can leverage syndecan-2 and potentially, other proteoglycans, to better support HSC self-renewal during homeostasis and upon stress.

## ACKNOWLEDGMENTS

We are grateful to Cyd McKay, Christina Root, Alex Hastie, and LaKeisha Perkins for technical assistance with animal studies. We would like to acknowledge the excellent assistance provided by the Fred Hutchinson Cancer Center Shared Resources: Comparative Medicine, Experimental Histopathology (Audrey Heintz and Stephanie Weaver), Flow Cytometry and Genomics. The investigation was supported in part by 5K01DK126989 (CMT), 23CDA1039196 from the American Heart Association (CMT), a New Investigator Award from the American Society for Transplantation and Cellular Therapy (CMT), a Fellowship from the American Society of Gene and Cell Therapy (CMT), a Distinguished Research Award from the Andy Hill CARE Foundation, and a V Scholar Award V2024-028 (CMT). KAW was supported by a Fellowship from the Fred Hutchinson Cancer Center Office of Faculty Affairs, the Graduate Research Fellowship Program from the National Science Foundation (DGE-2140004), and 1 R21 CA283589-01S1. CN was supported by a SPARK Fellowship from the Damon Runyon Cancer Research Foundation (SPK-04-24). This research was supported by NIH P30 CA015704 to the Fred Hutch/University of Washington/Seattle Children’s Cancer Consortium, which includes the Comparative Medicine, Experimental Histopathology and Flow Cytometry Shared Resources. We are grateful to Dr. John Chute for sharing the *Sdc2*^flox/flox^ strain with us and would like to thank Amara Pang for assistance generating this strain.

## AUTHOR CONTRIBUTIONS

Project conception and supervision (CMT), performed experiments (MWH, NJS, KAW, TMB, CN, MNN), analyzed data (RW, BKH, KAW, MWH, NJS, MNN, CMT), made figures (CMT, MWH, RW), wrote the manuscript with input from all authors (CMT, BKH, MWH).

## CONFLICT OF INTEREST

The authors declare no competing financial interests or conflicts of interest.

## DATA AND CODE SHARING STATEMENT

Original data are available to other investigators through reasonable requests submitted to the corresponding author. RNA sequencing data will be deposited in GEO when the government shutdown is resolved. Code is available at https://github.com/FredHutch/Hagen-et-al-2026.

## Notes

### Competing Interest Statement

The authors have declared no competing interest.

## References Cited

1. Acar, M. et al. (2015) Deep imaging of bone marrow shows non-dividing stem cells are mainly perisinusoidal. Nature 526, 126-+. 10.1038/nature15250

2. Arai, F. and Suda, T. (2007) Maintenance of quiescent hematopoietic stem cells in the osteoblastic niche. Ann N Y Acad Sci 1106, 41–53. 10.1196/annals.1392.005

3. Chen, Q. et al. (2019) Apelin(+) Endothelial Niche Cells Control Hematopoiesis and Mediate Vascular Regeneration after Myeloablative Injury. Cell Stem Cell 25, 768–783 e766. 10.1016/j.stem.2019.10.006

4. Ding, L. et al. (2012) Endothelial and perivascular cells maintain haematopoietic stem cells. Nature 481, 457–462. 10.1038/nature10783

5. Kunisaki, Y. et al. (2013) Arteriolar niches maintain haematopoietic stem cell quiescence. Nature 502, 637–643. 10.1038/nature12612

6. Szade, K. et al. (2018) Where Hematopoietic Stem Cells Live: The Bone Marrow Niche. Antioxid Redox Sign 29, 191–204. 10.1089/ars.2017.7419

7. Himburg, H.A. et al. (2017) Dickkopf-1 promotes hematopoietic regeneration via direct and niche-mediated mechanisms. Nat Med 23, 91–99. 10.1038/nm.4251

8. Himburg, H.A. et al. (2014) Pleiotrophin mediates hematopoietic regeneration via activation of RAS. J Clin Invest 124, 4753–4758. 10.1172/JCI76838

9. Termini, C.M. et al. (2021) Neuropilin 1 regulates bone marrow vascular regeneration and hematopoietic reconstitution. Nat Commun 12, 6990. 10.1038/s41467-021-27263-y

10. Xu, C. et al. (2018) Stem cell factor is selectively secreted by arterial endothelial cells in bone marrow. Nat Commun 9, 2449. 10.1038/s41467-018-04726-3

11. Himburg, H.A. et al. (2018) Distinct Bone Marrow Sources of Pleiotrophin Control Hematopoietic Stem Cell Maintenance and Regeneration. Cell Stem Cell 23, 370–381 e375. 10.1016/j.stem.2018.07.003

12. Zhou, B.O. et al. (2017) Bone marrow adipocytes promote the regeneration of stem cells and haematopoiesis by secreting SCF. Nat Cell Biol 19, 891–903. 10.1038/ncb3570

13. Sugiyama, T. et al. (2006) Maintenance of the hematopoietic stem cell pool by CXCL12-CXCR4 chemokine signaling in bone marrow stromal cell niches. Immunity 25, 977–988. 10.1016/j.immuni.2006.10.016

14. Renders, S. et al. (2021) Niche derived netrin-1 regulates hematopoietic stem cell dormancy via its receptor neogenin-1. Nat Commun 12, 608. 10.1038/s41467-020-20801-0

15. Balasubramanian, R. and Zhang, X. (2016) Mechanisms of FGF gradient formation during embryogenesis. Semin Cell Dev Biol 53, 94–100. 10.1016/j.semcdb.2015.10.004

16. Bishop, J.R. et al. (2007) Heparan sulphate proteoglycans fine-tune mammalian physiology. Nature 446, 1030–1037. 10.1038/nature05817

17. Matsuo, I. and Kimura-Yoshida, C. (2013) Extracellular modulation of Fibroblast Growth Factor signaling through heparan sulfate proteoglycans in mammalian development. Curr Opin Genet Dev 23, 399–407. 10.1016/j.gde.2013.02.004

18. McCaffrey, T.A. et al. (1992) Transforming growth factor-beta 1 is a heparin-binding protein: identification of putative heparin-binding regions and isolation of heparins with varying affinity for TGF-beta 1. J Cell Physiol 152, 430–440. 10.1002/jcp.1041520226

19. Lyon, M. et al. (1997) The interaction of the transforming growth factor-betas with heparin/heparan sulfate is isoform-specific. J Biol Chem 272, 18000–18006. 10.1074/jbc.272.29.18000

20. Murphy, J.W. et al. (2007) Structural and functional basis of CXCL12 (stromal cell-derived factor-1 alpha) binding to heparin. J Biol Chem 282, 10018–10027. 10.1074/jbc.M608796200

21. Kishimoto, S. et al. (2009) Human stem cell factor (SCF) is a heparin-binding cytokine. J Biochem 145, 275–278. 10.1093/jb/mvn169

22. Termini, C.M. et al. (2022) Syndecan-2 enriches for hematopoietic stem cells and regulates stem cell repopulating capacity. Blood 139, 188–204. 10.1182/blood.2020010447

23. Bruno, E. et al. (1995) Marrow-derived heparan sulfate proteoglycan mediates the adhesion of hematopoietic progenitor cells to cytokines. Exp Hematol 23, 1212–1217

24. Gupta, P. et al. (1998) Structurally specific heparan sulfates support primitive human hematopoiesis by formation of a multimolecular stem cell niche. Blood 92, 4641–4651

25. Schofield, K.P. et al. (1999) Expression of proteoglycan core proteins in human bone marrow stroma. Biochem J 343 Pt 3, 663–668

26. Siczkowski, M. et al. (1992) Binding of primitive hematopoietic progenitor cells to marrow stromal cells involves heparan sulfate. Blood 80, 912–919

27. Roberts, R. et al. (1988) Heparan sulphate bound growth factors: a mechanism for stromal cell mediated haemopoiesis. Nature 332, 376–378. 10.1038/332376a0

28. Amend, S.R. et al. (2016) Murine Hind Limb Long Bone Dissection and Bone Marrow Isolation. J Vis Exp. 10.3791/53936

29. Baryawno, N. et al. (2019) A Cellular Taxonomy of the Bone Marrow Stroma in Homeostasis and Leukemia. Cell 177, 1915–1932 e1916. 10.1016/j.cell.2019.04.040

30. Dolgalev, I. and Tikhonova, A.N. (2021) Connecting the Dots: Resolving the Bone Marrow Niche Heterogeneity. Front Cell Dev Biol 9, 622519. 10.3389/fcell.2021.622519

31. Kusumbe, A.P. et al. (2015) Sample preparation for high-resolution 3D confocal imaging of mouse skeletal tissue. Nat Protoc 10, 1904–1914. 10.1038/nprot.2015.125

32. Graf, L. et al. (2002) Gene expression profiling of the functionally distinct human bone marrow stromal cell lines HS-5 and HS-27a. Blood 100, 1509–1511. 10.1182/blood-2002-03-0844

33. Ogawa, M. et al. (1997) In vitro expansion of hematopoietic stem cells. Stem Cells 15 Suppl 1, 7–11; discussion 12. 10.1002/stem.5530150803

34. Traycoff, C.M. et al. (1996) Ex vivo expansion of murine hematopoietic progenitor cells generates classes of expanded cells possessing different levels of bone marrow repopulating potential. Exp Hematol 24, 299–306

35. van der Loo, J.C. and Ploemacher, R.E. (1995) Marrow- and spleen-seeding efficiencies of all murine hematopoietic stem cell subsets are decreased by preincubation with hematopoietic growth factors. Blood 85, 2598–2606

36. Wilkinson, A.C. et al. (2020) Haematopoietic stem cell self-renewal in vivo and ex vivo. Nat Rev Genet 21, 541–554. 10.1038/s41576-020-0241-0

37. Desterke, C. et al. (2021) EGR1 dysregulation defines an inflammatory and leukemic program in cell trajectory of human-aged hematopoietic stem cells (HSC). Stem Cell Res Ther 12, 419. 10.1186/s13287-021-02498-0

38. Browaeys, R. et al. (2020) NicheNet: modeling intercellular communication by linking ligands to target genes. Nat Methods 17, 159–162. 10.1038/s41592-019-0667-5

39. Piszczatowski, R.T. et al. (2024) Heparan Sulfates and Heparan Sulfate Proteoglycans in Hematopoiesis. Blood. 10.1182/blood.2023022736

40. Crowley, S.J. et al. (2025) BCLAF1 restrains stress responses in hematopoietic stem cells to support expansion and repopulation. Blood Adv. 10.1182/bloodadvances.2024014916

41. Okada, S. et al. (1999) Prolonged expression of c-fos suppresses cell cycle entry of dormant hematopoietic stem cells. Blood 93, 816–825

42. Xiao, X. et al. (2019) Targeting JNK pathway promotes human hematopoietic stem cell expansion. Cell Discov 5, 2. 10.1038/s41421-018-0072-8

43. Oedekoven, C.A. et al. (2021) Hematopoietic stem cells retain functional potential and molecular identity in hibernation cultures. Stem Cell Reports 16, 1614–1628. 10.1016/j.stemcr.2021.04.002

44. Comazzetto, S. et al. (2025) Ascorbate deficiency increases quiescence and self-renewal in hematopoietic stem cells and multipotent progenitors. Blood 145, 114–126. 10.1182/blood.2024024769

45. Huang, B. et al. (2008) SCF-mediated mast cell infiltration and activation exacerbate the inflammation and immunosuppression in tumor microenvironment. Blood 112, 1269–1279. 10.1182/blood-2008-03-147033

46. Sundqvist, A. et al. (2020) TGFbeta and EGF signaling orchestrates the AP-1- and p63 transcriptional regulation of breast cancer invasiveness. Oncogene 39, 4436–4449. 10.1038/s41388-020-1299-z

47. Yamazaki, S. et al. (2009) TGF-beta as a candidate bone marrow niche signal to induce hematopoietic stem cell hibernation. Blood 113, 1250–1256. 10.1182/blood-2008-04-146480

48. Ding, L. and Morrison, S.J. (2013) Haematopoietic stem cells and early lymphoid progenitors occupy distinct bone marrow niches. Nature 495, 231–235. 10.1038/nature11885

49. Yoshihara, H. et al. (2007) Thrombopoietin/MPL signaling regulates hematopoietic stem cell quiescence and interaction with the osteoblastic niche. Cell Stem Cell 1, 685–697. 10.1016/j.stem.2007.10.020

50. Qian, H. et al. (2007) Critical role of thrombopoietin in maintaining adult quiescent hematopoietic stem cells. Cell Stem Cell 1, 671–684. 10.1016/j.stem.2007.10.008

