## Supplemental Materials for "Depletion of microenvironmental syndecan-2 impairs hematopoietic stem cell self-renewal and cytokine responses"

**SUPPLEMENTAL METHODS**

**Generation of transgenic mice:** We generated *Sdc2*^flox/flox^ mice by purchasing frozen sperm from the European Mouse Mutant Association (EMMA) for C57BL/7NTac-Sdc2tm1a(KOMP)Wtsi/Ieg (EMMA ID 06284). The Jackson Laboratory (Bar Harbor, ME) received these sperm from EMMA, which were thawed and used in assisted IVF using stock# 5304 C57BL/6NJ oocytes. Embryos were transferred for a maximum number of 4-5 embryo transfers (~15 embryos per transfer). The resulting heterozygous male mice from the cryorecovery were bred with Homozygous B6.129S4-Gt(ROSA)26Sortm1(FLP1)Dym/RainJ (Jax Strain 009086) females. The resulting mice were bred to produce homozygous mice for us to use to breed with Cre-recombinase mice.

*Sdc2^flox/flox^* mice were crossed with the following tissue-specific Cre-recombinase mice in the B57BL/6 background purchased from Jackson Laboratories: *Vav1-Cre*, *Cdh5-Cre*, and *Lepr-Cre* (see resources table). Cre-recombinase mice were crossed with Sdc2flox/flox mice and their hemizygous offspring (Cre+/-; *Sdc2*^flox/-^) were crossed to generate either *Cre-*; *Sdc2*^flox/flox^ or *Cre+*; *Sdc2*^floxflox^ mice, which were used in downstream experimental applications. Endothelial and stromal reporters were generated by crossing tissue-specific *Cdh5-cre* or *Lepr-cre* mice with a Rosa26 tdTomato reporter; F1 mice are used for experiments.

**Confocal imaging:** Mice were infused with anti-VE-cadherin antibody and femurs were extracted and prepared, as described (1, 2). Sections were stained with anti-syndecan-2 antibody (1 µg/sample), washed, and stained with anti-sheep alexa-488 (1:200) and DAPI (1:1000). Slides were mounted with Fluoromount G and imaged using a Leica SP8 system with a 20X objective.

**Conventional histology:** One femur per mouse was fixed in formalin for 48-72 hours, washed in PBS, and decalcified (0.5M EDTA pH 7.4) for 14 days at 4ºC with constant nutation. Decalcified femurs were paraffin embedded, sectioned at 4µm, and stained with hematoxylin and eosin by the Fred Hutch Experimental Histopathology Shared Resource. Full-section scans were generated using a digital pathology scanner with a 20X objective. Images were analyzed by creating serial random forest classifiers using HALO. Broad regions of interest encompassing the BM compartment, excluding the epiphysis, were defined by a single observer. Primary classifiers were trained to segregate the central marrow compartment from glass, cortical bone, and fold and tear artifacts. Secondary classifiers identified cellular, acellular, or RBC area. Area fractions (positively classified area divided by all marrow area) are reported.

**Reciprocal chimeras:** 1x10^6^ whole bone marrow cells were intravenously injected into Control or *Sdc2*^∆MSC^ mice (10Gy, ^137^Cs), which achieves >90% donor cell chimerism in the peripheral blood. Peripheral blood and bone marrow were analyzed, as described for competitive repopulation assays.

**ShRNA transfection:** Non-treated tissue culture treated plastic was coated with Retronectin (50µg mL^-1^) and incubated at room temperature for 2 hours followed by 30 minutes with bovine serum albumin (2% v/v in PBS). 5×10^4^ of BMEC-1 or HS-5 cells were rapidly thawed and plated. Lentiviruses were added to a multiplicity of infection of 10 and centrifuged for 30 minutes at 235xg at 32ºC. Transduced HS-5 cells were maintained in complete Advanced DMEM (Advanced DMEM supplemented with 10% FBS and 1% penicillin-streptomycin) and transduced BMEC-1 cells were maintained in complete MCDB-131 (MCDB-131 supplemented with 10% FBS and 1% penicillin-streptomycin), which was refreshed every 48 hours.

**scRNA sequencing analysis:** FASTQ files were processed with Cell Ranger v9.0.0 to align reads to the mm10 version 2020-A reference genome. Individual samples were filtered for cells with 500 < unique molecular identifiers (UMIs) < 100,000 and percent mitochondrial transcripts < 5%. Doublets were removed using Scrublet. Using Seurat, UMI counts were normalized with default inputs, variable features were set to the maximum number of detected features for each sample, and the data was scaled using all features. Data were dimensionally reduced using 50 principle components and 2-dimensional unique manifold approximation projection (UMAP). The Seurat object was then converted into a cell dataset object for use with Monocle 3. Individual cells were annotated with cell types using Garnett with default outgroup cell settings and the cell type markers from Itkin et al. 2023 listed in Supplementary Table 2. Clusters were identified with Monocle 3 using a resolution of 5e-4. Clusters were classified as the cell type maximally represented with a couple of exceptions: one of the clusters annotated with “HSC” was reclassified as “MPP”, which was almost equally represented; and several clusters were reclassified from “unknown” to the next most represented cell type. Differentially expressed genes (DEGs) between conditions in HSCs were identified using a regression fit to a quasipoisson model in Monocle 3. DEGs were filtered for genes with q-values < 0.05 and -0.25 ≥ log_2_(fold change) ≥ 0.25. This list of DEGs was utilized in NicheNetR to identify ligand-receptor signaling associated with the differential expression profile.

**Surface plasmon resonance:** SPR analysis was performed as a service by MedChemExpress (Monmouth Junction, NJ) using CM5 sensor chips and amine coupling kit from Cytiva and a Biacore instrument from GE Healthcare.

**Heparan sulfate binding assay:** Biotinylation and the heparan sulfate binding assay were performed according to the protocol, as described (3). Human TGF-β1 was biotinylated using Heparin Sepharose 6 Fast Flow Resin and EZ-link Sulfo-NHS-LC-Biotin. 1-2.5 x 10^5^ shControl or sh*SDC2* cells were seeded in advanced DMEM media and incubated overnight at 37°C, 5% CO_2_. Cells were dissociated using a 10 mM EDTA-PBS solution and incubated with 50 ng of biotin-TGF-β1 for 1 hour on ice. Unbound TGF-β1 was washed away with 0.1% BSA in PBS. Cells were then stained with Streptavidin-AlexaFluor647 for 20 minutes and analyzed via flow cytometry. AlexaFluor647 MFI was calculated for each sample.

**SUPPLEMENTAL FIGURE LGENDS**

**Supplemental Figure 1: Generation and characterization of *Sdc2* transgenic mice.** **(A)** Schematic depicting the *Sdc2* transgenic allele in the C57BL/7NTac-Sdc2tm1a(KOMP)Wtsi/Ieg strain generated by the European Mouse Mutant Association, which was cryoderived and crossed according to the depicted schemes to generate the genetic lines used in this study. Flow cytometric analysis of syndecan-2 surface expression on **(B)** CD45^-^Leptin Receptor^+^ bone marrow mesenchymal stromal cells from adult Control or *Sdc2*^∆MSC^ mice, (C) CD45^-^VE-Cadherin bone marrow endothelial cells from adult Control or *Sdc2*^∆EC^ mice, or (D) CD45^+^ bone marrow hematopoietic cells from Control or *Sdc2*^∆HC^ mice. (*n=3-7 biological replicates/strain, statistics show unpaired t-tests*).

**Supplemental Figure 2: Bone marrow analysis of *Sdc2* transgenic mice.** Micrographs depicting femur bone marrow sections from **(A)** Control or *Sdc2*^∆MSC^ mice, **(B)** Control or *Sdc2*^∆EC^ mice, or **(C)** Control or *Sdc2*^∆HC^ mice stained with hematoxylin and eosin. Bone marrow cellularity quantification from femurs from **(D)** Control or *Sdc2*^∆MSC^ mice, **(E)** Control or *Sdc2*^∆EC^ mice, or **(F)** Control or *Sdc2*^∆HC^ mice. The frequency of bone marrow B220^+^ B cells, CD3^+^ T cells, and Mac-1/Gr-1^+^ myeloid cells was quantified in femurs from **(G)** Control or *Sdc2*^∆MSC^ mice, **(I)** Control or *Sdc2*^∆EC^ mice, or **(I)** Control or *Sdc2*^∆HC^ mice. Representative flow cytometric gating of bone marrow hematopoietic progenitor cells and quantification in **(J)** Control or *Sdc2*^∆MSC^ mice, **(K)** Control or *Sdc2*^∆EC^ mice, or **(L)** Control or *Sdc2*^∆HC^ mice. Representative flow cytometric gating of bone marrow hematopoietic stem and progenitor cells and quantification of hematopoietic stem and progenitor cells in the bone marrow of **(M)** Control or *Sdc2*^∆MSC^ mice, **(N)** Control or *Sdc2*^∆EC^ mice, or **(O)** Control or *Sdc2*^∆HC^ mice. *(n=3-6 biological replicates/strain, statistics show unpaired t-tests. *p<0.05, ***p<0.001).*

**Supplemental Figure 3: Bone marrow analysis of recipient mice after primary and secondary transplantation of cells from *Sdc2* transgenic mice.** Percent donor chimerism in the bone marrow overall and within the B220+ B cell, CD3+ T cell, and Mac-1/Gr-1 myeloid cell populations after primary competitive transplant with **(A)** Control or *Sdc2*^∆MSC^ cells, **(B)** Control or *Sdc2*^∆EC^ cells, or **(C)** Control or *Sdc2*^∆HC^ cells. Percent donor chimerism in the bone marrow overall and within the B220+ B cell, CD3+ T cell, and Mac-1/Gr-1 myeloid cell populations after secondary competitive transplant with **(D)** Control or *Sdc2*^∆MSC^ cells, **(E)** Control or *Sdc2*^∆EC^ cells, or **(F)** Control or *Sdc2*^∆HC^ cells. (*for primary transplants, n=10 recipients/strain; for secondary transplants, n=7-10 recipients per strain; statistics denote unpaired t-tests)*.

**Supplemental Figure 4: Hematopoietic cell transplant into microenvironments lacking Sdc2 in Lepr-targeted cells. (A)** Experimental design for hematopoietic cell transplant assay using bone marrow cells from SJL mice transplanted into lethally irradiated Control or *Sdc2*^∆MSC^ mice. **(B)** Complete blood count analysis at 24-weeks after bone marrow transplant into Control or *Sdc2*^∆MSC^ mice. **(C)** Percent B220^+^ B cells, CD3^+^ T cells, or Mac-1/Gr-1^+^ myeloid cells within donor cells in the peripheral blood after bone marrow transplant into Control or *Sdc2*^∆MSC^ mice. **(D)** Percent B220^+^ B cells, CD3^+^ T cells, or Mac-1/Gr-1^+^ myeloid cells within donor cells in the bone marrow at 24-weeks post bone marrow transplant into Control or *Sdc2*^∆MSC^ mice. **(E)** The frequency of hematopoietic stem and progenitor cells within donor cells in the bone marrow after bone marrow transplant into Control or *Sdc2*^∆MSC^ mice. (*n=4 recipients/strain, statistics show unpaired t-tests*).

**Supplementary Table 1: Manuscript Resources table.** Vendor information for antibodies, assays, mouse models, and software used throughout manuscript.

| **ANTIBODIES** | | | | | | | | | | |
| --- | --- | --- | --- | --- | --- | --- | --- | --- | --- | --- |
| **Name** | | | **Vendor** | | | **Catalog Number** | **Clone** | | | **Dilution** |
| V450 Mouse Lineage Antibody Cocktail with isotype | | | BD Biosciences | | | 561301 | n/a | | | 20µL / 10^6^ cells |
| CD45Cyanine 7 anti-mouse Ly-6A/E | | | Biolegend | | | 108125 | D7 | | | 1µg / 10^6^ cells |
| APC/Cyanine 7 Rat IgG2a, κ Isotype control | | | Biolegend | | | 400523 | RTK2758 | | | 1µg / 10^6^ cells |
| PE anti-mouse CD117 | | | Biolegend | | | 105807 | 2B8 | | | 1µg / 10^6^ cells |
| PE Rat IgG2b, κ Isotype control | | | Biolegend | | | 400607 | RTK4530 | | | 1µg / 10^6^ cells |
| PE/Cyanine 7 anti-mouse CD34 | | | Biolegend | | | 119325 | MEC14.7 | | | 1µg / 10^6^ cells |
| PE/Cyanine 7 Rat IgG2a, κ Isotype control | | | Biolegend | | | 400521 | RTK2758 | | | 1µg / 10^6^ cells |
| KIRAVIA Blue 520 anti-mouse CD135 | | | Biolegend | | | 135323 | A2F10 | | | 1µg / 10^6^ cells |
| KIRAVIA Blue 520 Rat IgG2a, κ isotype control | | | Biolegend | | | 400575 | RTK2758 | | | 1µg / 10^6^ cells |
| Brilliant Violet 605 anti-mouse CD150 | | | Biolegend | | | 115927 | TC15-12F12.2 | | | 1µg / 10^6^ cells |
| Brilliant Violet 605 Rat IgG2a, κ isotype control | | | Biolegend | | | 400539 | RTK2758 | | | 1µg / 10^6^ cells |
| Alexa Fluor 700 anti-mouse CD48 | | | Biolegend | | | 103425 | HM48-1 | | | 1µg / 10^6^ cells |
| Hamster IgG Isotype Ctrl | | | Biolegend | | | 400926 | HTK888 | | | 1µg / 10^6^ cells |
| APC Rat Anti-Mouse TER-119 / Erythroid cells | | | BD Biosciences | | | 557909 | TER-119 | | | 0.4µg / 10^5^ cells |
| APC Rat IgG2b, κ isotype control | | | BD Biosciences | | | 553991 |  | | | 1µg / 10^6^ cells |
| Brilliant Violet 421 anti-mouse CD3 | | | Biolegend | | | 100228 | 17A2 | | | 0.4µg / 10^5^ cells |
| Brilliant Violet 421 Rat IgG2b, κ Isotype Control | | | Biolegend | | | 400639 | RTK4530 | | | 1µg / 10^6^ cells |
| BV711 Rat Anti-Mouse CD45R/B220 | | | BD Biosciences | | | 563892 | RA3-6B2 | | | 0.4µg / 10^5^ cells |
| BV711 Hamster IgG1, κ isotype control | | | BD Biosciences | | | 563128 |  | | | 1µg / 10^6^ cells |
| PE Rat Anti-CD11b | | | BD Biosciences | | | 553311 | M1/70 | | | 0.4µg / 10^5^ cells |
| PE Rat Anti-Mouse Ly-6G and Ly-6C | | | BD Biosciences | | | 553128 | RB6-6C5 | | | 0.4µg / 10^5^ cells |
| PE Rat IgG2b, κ Isotype control | | | BD Biosciences | | | 553989 |  | | | 1µg / 10^6^ cells |
| Purified Rat Anti-Mouse CD16/CD32 (Mouse BD FC Block) | | | BD Biosciences | | | 553142 | 2.4G2 | | | 2.5µg / 10^6^ cells |
| Alexa Fluor 700 Rat Anti-Mouse CD45 | | | Biolegend | | | 103128 | 30-F11 | | | 1µg / 10^6^ cells |
| Alexa Fluor 700 Rat IgG2b, κ isotype control | | | Biolegend | | | 400628 | RTK4530 | | | 1µg / 10^6^ cells |
| CD31 | | | Cell signaling technologies | | | 77699S |  | | | 1:200 |
| APC Rat Anti-Mouse TER-119 / Erythroid cells | | | Biolegend | | | 116212 | TER-119 | | | 1µg / 10^6^ cells |
| APC Rat IgG2b, κ Isotype Ctrl Antibody | | | Biolegend | | | 400612 | RTK4530 | | | 1µg / 10^6^ cells |
| Brilliant Violet 711™ anti-mouse/human CD45R/B220 Antibody | | | Biolegend | | | 103255 | RA3-6B2 | | | 1µg / 10^6^ cells |
| Brilliant Violet 711™ Rat IgG2a, κ Isotype Ctrl Antibody | | | Biolegend | | | 400551 | RTK2758 | | | 1µg / 10^6^ cells |
| PE anti-mouse/human CD11b Antibody | | | Biolegend | | | 101208 | M1/70 | | | 1µg / 10^6^ cells |
| PE Rat IgG2b, κ Isotype Ctrl Antibody | | | Biolegend | | | 400636 | RTK4530 | | | 1µg / 10^6^ cells |
| PE anti-mouse Ly-6G/Ly-6C (Gr-1) Antibody | | | Biolegend | | | 108408 | RB6-8C5 | | | 1µg / 10^6^ cells |
| Brilliant Violet 605™ anti-mouse CD45.1 Antibody | | | Biolegend | | | 110737 | A20 | | | 1µg / 10^6^ cells |
| Brilliant Violet 605™ Mouse IgG2a, κ Isotype Ctrl Antibody | | | Biolegend | | | 400270 | MOPC-173 | | | 1µg / 10^6^ cells |
| Alexa Fluor® 700 anti-mouse 4002 Antibody | | | Biolegend | | | 109822 | 104 | | | 1µg / 10^6^ cells |
| APC anti-mouse CD45.2 Antibody | | | Biolegend | | | 109813 | 104 | | | 1µg / 10^6^ cells |
| APC Mouse IgG2a, κ Isotype Ctrl (FC) Antibody | | | Biolegend | | | 400221 | 17A2 | | | 1µg / 10^6^ cells |
| Alexa Fluor® 700 Mouse IgG2a, κ Isotype Ctrl Antibody | | | Biolegend | | | 400248 | MOPC-173 | | | 1µg / 10^6^ cells |
| BD Pharmingen™ FITC Mouse Anti-Mouse CD45.1 | | | BD Biosciences | | | 553775 | A20 | | | 1µg / 10^6^ cells |
| PE/Cyanine7 anti-mouse CD45.1 Antibody | | | Biolegend | | | 110730 | A20 | | | 1µg / 10^6^ cells |
| PE/Cyanine7 Mouse IgG2a, κ Isotype Ctrl Antibody | | | Biolegend | | | 400253 | MOPC-173 | | | 1µg / 10^6^ cells |
| Brilliant Violet 605™ anti-mouse Ly-6A/E (Sca-1) Antibody | | | Biolegend | | | 108134 | D7 | | | 1µg / 10^6^ cells |
| BD Horizon™ BV605 Mouse IgG2a, k Isotype Control | | | BD Biosciences | | | 562778 | G155-178 | | | 1µg / 10^6^ cells |
| PE anti-mouse CD34 Antibody | | | Biolegend | | | 119308 | MEC14.7 | | | 1µg / 10^6^ cells |
| BD Pharmingen™ PE Rat IgG2a, κ Isotype Control | | | BD Biosciences | | | 553930 | R35-95 | | | 1µg / 10^6^ cells |
| BD Pharmingen™ APC-H7 Rat anti-Mouse CD117 | | | BD Biosciences | | | 560185 | 2B8 | | | 1µg / 10^6^ cells |
| BD Pharmingen™ APC-H7 Rat IgG2b, κ Isotype Control | | | BD Biosciences | | | 560200 | A95-1 | | | 1µg / 10^6^ cells |
| Alexa Fluor® 647 anti-mouse CD144 (VE-cadherin) Antibody | | | Biolegend | | | 138006 | BV13 | | |  |
| **MOUSE MODELS** | | | | | | | | | | |
| **Name** | | | **Vendor** | | | | **Catalog Number** | | | |
| Mouse C57BL/6J | | | The Jackson Laboratory | | | | 000664 | | | |
| Mouse CDH5 Cre (B6.FVB-Tg(Cdh5-cre)7Mlia/J) | | | The Jackson Laboratory | | | | 6137 | | | |
| Mouse TdTomato (B6.Cg-Gt(ROSA)26Sortm14(CAG-tdTomato)Hze/J) | | | The Jackson Laboratory | | | | 7914 | | | |
| Mouse Lepr Cre (B6.129(Cg)-Leprtm2(cre)Rck/J) | | | The Jackson Laboratory | | | | 008320 | | | |
| Mouse Vav-iCre (B6.Cg-Commd10Tg(Vav1-icre)A2Kio/J) | | | The Jackson Laboratory | | | | 008610 | | | |
| **CELL LINES** | | | | | | | | | | |
| **Name** | | | | **Vendor** | | | **Catalog number** | | | |
| HS-5 | | | | ATCC | | | CRL-3611 | | | |
| BMEC-1 | | | | ATCC | | | CRL-3421 | | | |
| **LENTIVIRAL VECTORS** | | | | | | | | | | |
| **Name** | **Titer** | | | | **Vendor** | | | | **Catalog number** | |
| pLKO.1_U6-shRNA:hPGK-Puro-CMV (SHC016 Negative Ctrl) | 1.2x10^9^ VP/mL | | | | Millipore Sigma | | | | 09172429MN | |
| TRCN0000123092 pLKO.1-puro-CMV-tGFP CSTVRS SDC2 Human) | 5.1x10^7^ VP/mL | | | | Millipore Sigma | | | | 08222305MN | |
| **qPCR PROBES** | | | | | | | | | | |
| **Name** | | | **Vendor** | | | | **Catalog number** | | | |
| Mm00448918_m1 (Mouse Sdc1) | | | Thermo Fisher Scientific | | | | 4331182 | | | |
| Mm04207492_m1 (Mouse Sdc2) | | | Thermo Fisher Scientific | | | | 4331182 | | | |
| Mm01179833_m1 (Mouse Sdc3) | | | Thermo Fisher Scientific | | | | 4331182 | | | |
| Mm00488527_m1 (Mouse Sdc4) | | | Thermo Fisher Scientific | | | | 4331182 | | | |
| Hs04966523_m1 (Human SDC1) | | | Thermo Fisher Scientific | | | | 4331182 | | | |
| Hs01081432_m1 (Human SDC2) | | | Thermo Fisher Scientific | | | | 4331182 | | | |
| Hs01568665_m1 (Human SDC3) | | | Thermo Fisher Scientific | | | | 4331182 | | | |
| Hs01120908_m1 (Human SDC4) | | | Thermo Fisher Scientific | | | | 4331182 | | | |
| **REAGENTS** | | | | | | | | | | |
| **Name** | | | **Vendor** | | | | **Catalog number** | | | |
| Iscove's Modified Dulbecco's Medium | | | Gibco | | | | 12440-053 | | | |
| Penicillin-Streptomycin Solution | | | Fisher Scientific | | | | MT30002CI | | | |
| Fetal bovine serum | | | Fisher Scientific | | | | 501527078 | | | |
| Compensation beads | | | Fisher Scientific | | | | 3031-5813-28 | | | |
| 7AAD cell viability stain | | | BD Biosciences | | | | 559925 | | | |
| Methocult GF M3434 | | | Stemcell Technologies | | | | 03434 | | | |
| SmartDish | | | Stemcell Technologies | | | | 27371 | | | |
| EDTA vacutainers | | | BD | | | | 101005 | | | |
| Ammonium-Chloride-Potassium (ACK) Lysing Buffer | | | Fisher Scientific | | | | 50-983-219 | | | |
| Applied Biosystems™ TaqMan™ Universal PCR Master Mix | | | Fisher Scientific | | | | 43-044-37 | | | |
| Liberase | | | Millipore Sigma | | | | 5401020001 | | | |
| Bovine Serum Albumin | | | Sigma Aldrich | | | | A7030-100G | | | |
| RetroNectin® Recombinant Human Fibronectin Fragment | | | TakaraBio | | | | T100A | | | |
| Human TGF-β1 | | | Stemcell Technologies | | | | 78067.1 | | | |
| **COMMERCIAL KITS** | | | | | | | | | | |
| **Name** | | | **Vendor** | | | | **Catalog Number** | | | |
| Direct lineage cell depletion kit mouse | | | Miltenyi | | | | 130-110-470 | | | |
| LS columns | | | Miltenyi | | | | 130-042-401 | | | |
| RNeasy Micro Kit | | | Qiagen | | | | 74004 | | | |
| **INSTRUMENTS** | | | | | | | | | | |
| **Model** | | **Vendor** | | | | | | **Purpose** | | |
| LSRFortessa X-50 | | BD Biosciences | | | | | | Flow cytometry | | |
| FACSymphony A5 | | BD Biosciences | | | | | | Flow cytometry | | |
| FACSymphony S6 | | BD Biosciences | | | | | | FACS & flow cytometry | | |
| TC-20 | | Bio-Rad | | | | | | Automated cell counting | | |
| Mastercycler Nexus X2 | | Eppendorf | | | | | | PCR | | |
| QuantStudio5 | | Applied Biosystems | | | | | | qRT-PCR | | |
| Element HT5 | | Antech | | | | | | Complete blood count | | |
| SP8 | | Leica | | | | | | Confocal microscopy | | |
| Aperio Versa200 | | Leica | | | | | | Brightfield microscopy | | |
| Chromium controller | | 10X Genomics | | | | | | scRNA library preparation | | |
| NovaSeq X+ 10B | | Illumina | | | | | | RNA Sequencing | | |
| **SOFTWARE** | | | | | | | | | | |
| **Name** | | | **Vendor/Publication** | | | | **Details** | | | |
| HALO v.3.6.4134.265 | | | Indica Labs | | | |  | | | |
| FlowJo v10 | | | BD Biosciences | | | |  | | | |
| GraphPad Prism Version 10.4.1 (532) | | | GraphPad Software | | | |  | | | |
| STEMVision Analyzer v2.6.7.0 | | | Stemcell Technologies | | | |  | | | |
| STEMVision Colony Marker v2.5.0.0 | | | Stemcell Technologies | | | |  | | | |
| R v4.4.2 | | | R foundation for statistical computing | | | |  | | | |
| Cell Ranger v9.0.0 | | | 10X Genomics, Zheng et al. 2017 (8) | | | | Zheng GXY, Terry JM, Belgrader P, et al. Massively parallel digital transcriptional profiling of single cells. Nat Commun. 2017;8: 14049. doi: 10.1038/ncomms14049 | | | |
| Scrublet v.0.2.3 | | | Wolock et al. 2019 (9) | | | | Wolock SL, Lopez R, Klein AM. Scrublet: computational identification of cell doublets in single-cell transcriptomic data. Cell Syst. 2019;8(4):281-291. doi: 10.1016/j.cels.2018.11.005 | | | |
| Seurat v.5.2.1 | | | Hao et al. 2023 (10) | | | | Hao Y, Stuart T, Kowalski MH, et al. Dictionary learning for integrative, multimodal and scalable single-cell analysis. Nat Biotechnol. 2024;42(2):293-304. doi: 10.1038/s41587-023-01767-y | | | |
| Monocle 3 v.1.3.7 | | | Cao et al. 2019 (11) | | | | Cao J, Spielmann M, Qiu X, et al. The single-cell transcriptional landscape of mammalian organogenesis. Nature. 2019; 566(7745):496-502. doi: 10.1038/s41586-019-0969-x | | | |
| Garnett v.0.2.22 | | | Pliner et al. 2019 (12) | | | | Pliner HA, Shendure J, Trapnell C. Supervised classification enables rapid annotation of cell atlases. Nat Methods. 2019; 16(10):983-986. doi: 10.1038/s41592-019-0535-3 | | | |
| NicheNetR v.2.2.0 | | | Browaeys et al. 2020 (13) | | | | Browaeys R, Saelens W, Saeys Y. NicheNet: modeling intercellular communication by linking ligands to target genes. Nat Methods. 2020; 17(2):159-162. doi: 10.1038/s41592-019-0667-5 | | | |

**Supplementary Table 2: Garnett Markers.** Markers from Itkins et al. 2023 used for cell type classification via Garnett.

| **Cell Type** | **Expressed** | **Not Expressed** |
| --- | --- | --- |
| HSC | Hoxb5, Fgd5, Cdkn1c, Procr | Cd48, Flt3, Cd34 |
| MPP | CD34, Hlf | Hoxb5, Cdkn1c, Fgd5, Cebpe, Fcnb, Elane, Klf1, Gp1bb, Gata1, Csf1r, Siglech, Fcer1a, Il2rb, Procr, Vwf, Dntt, Rag2 |
| LPP | Dntt, Rag2 | Hlf, Procr, Gata2, Siglech, Ccl5, Il2rb, Fcer1a, Ms4a2 |
| MEP | Gata1, Zfpm1 | Fcer1a, Ms4a2, Procr, Fgd5, Hoxb5, Ly6a, Flt3, Ccl5, Il2rb |
| Monocyte | Adgre1, Csf1r | Siglech, Fcnb |
| Neutrophil | Fcnb, Elane, Cebpe | Adgre1, Ly86, Siglech |
| Dendritic | Siglech | Adgre1, Fcnb, Elane |
| Mast-Basophil | Fcer1a, Ms4a2 | Pf4, Klf1 |
| Lymphocyte | Ccl5, Il2rb | Siglech, Csf1r |

**SUPPLEMENTAL CITATIONS**

1.	Termini CM, Pang A, Fang T, Roos M, Chang VY, Zhang Y, et al. Neuropilin 1 regulates bone marrow vascular regeneration and hematopoietic reconstitution. Nat Commun. 2021;12(1):6990.

2.	Kusumbe AP, Ramasamy SK, Starsichova A, Adams RH. Sample preparation for high-resolution 3D confocal imaging of mouse skeletal tissue. Nat Protoc. 2015;10(12):1904-14.

3.	Basu A, Weiss RJ. Glycosaminoglycan Analysis: Purification, Structural Profiling, and GAG-Protein Interactions. Methods Mol Biol. 2023;2597:159-76.

4.	Dobin A, Davis CA, Schlesinger F, Drenkow J, Zaleski C, Jha S, et al. STAR: ultrafast universal RNA-seq aligner. Bioinformatics. 2013;29(1):15-21.

5.	DeLuca DS, Levin JZ, Sivachenko A, Fennell T, Nazaire MD, Williams C, et al. RNA-SeQC: RNA-seq metrics for quality control and process optimization. Bioinformatics. 2012;28(11):1530-2.

6.	Wang L, Wang S, Li W. RSeQC: quality control of RNA-seq experiments. Bioinformatics. 2012;28(16):2184-5.

7.	Robinson MD, McCarthy DJ, Smyth GK. edgeR: a Bioconductor package for differential expression analysis of digital gene expression data. Bioinformatics. 2010;26(1):139-40.

8.	Zheng GX, Terry JM, Belgrader P, Ryvkin P, Bent ZW, Wilson R, et al. Massively parallel digital transcriptional profiling of single cells. Nat Commun. 2017;8:14049.

9.	Wolock SL, Lopez R, Klein AM. Scrublet: Computational Identification of Cell Doublets in Single-Cell Transcriptomic Data. Cell Syst. 2019;8(4):281-91 e9.

10.	Hao Y, Stuart T, Kowalski MH, Choudhary S, Hoffman P, Hartman A, et al. Dictionary learning for integrative, multimodal and scalable single-cell analysis. Nat Biotechnol. 2024;42(2):293-304.

11.	Cao J, Spielmann M, Qiu X, Huang X, Ibrahim DM, Hill AJ, et al. The single-cell transcriptional landscape of mammalian organogenesis. Nature. 2019;566(7745):496-502.

12.	Pliner HA, Shendure J, Trapnell C. Supervised classification enables rapid annotation of cell atlases. Nat Methods. 2019;16(10):983-6.

13.	Browaeys R, Saelens W, Saeys Y. NicheNet: modeling intercellular communication by linking ligands to target genes. Nat Methods. 2020;17(2):159-62.
